# A Versatile Deep Graph Contrastive Learning Framework for Single-cell Proteomics Embedding

**DOI:** 10.1101/2022.12.14.520366

**Authors:** Wei Li, Fan Yang, Fang Wang, Yu Rong, Bingzhe Wu, Han Zhang, Jianhua Yao

## Abstract

The advance of single-cell proteomics sequencing technology sheds light on the research in revealing the protein-protein interactions, the post-translational modifications, and the proteoform dynamics of proteins in a cell. However, the uncertainty estimation for peptide quantification, data missingness, severe batch effects and high noise hinder the analysis of single-cell proteomic data. It is a significant challenge to solve this set of tangled problems together, where existing methods tailored for single-cell transcriptome do not address. Here, we proposed a novel versatile framework scPROTEIN, composed of peptide uncertainty estimation based on a multi-task heteroscedastic regression model and cell embedding learning based on graph contrastive learning designed for single-cell proteomic data analysis. scPROTEIN estimated the uncertainty of peptide quantification, denoised the protein data, removed batch effects and encoded single-cell proteomic-specific embeddings in a unified framework. We demonstrate that our method is efficient for cell clustering, batch correction, cell-type annotation and clinical analysis. Furthermore, our method can be easily plugged into single-cell resolved spatial proteomic data, laying the foundation for encoding spatial proteomic data for tumor microenvironment analysis.

## Introduction

In the past, the application of single-cell sequencing technologies mainly focused on detecting the level of single-cell transcriptome^1^, while recent single-cell proteomic technologies are also advancing^2–13^. Although the mRNA level of genes was considered as a proxy of the protein level in the past decades^14,15^, the transcriptional level of genes and the content of proteins that ultimately perform functions have low consistency, indicating that single-cell transcriptome alone is insufficient to derive cellular protein levels^16,17^. Proteins are the ultimate executors of cellular function, and proteomics provides knowledge on deciphering cellular mechanisms through enzyme activity and post-translational modifications^10^. Single-cell proteomics enables the identification of activated pathways in therapy-resistant cells and therefore provides biomarkers for cancer diagnosis and prognosis^18^. In recent years, improvements in the sensitivity of mass spectrometry and innovations in single-cell proteomic technologies have made it possible to quantify a range of proteins at the single-cell resolution^4,13^.

Despite the above advantages, single-cell proteomic data have several serious problems such as uncertainty of peptides quantification, data missingness, batch effects and data noise due to the limitations in current sample preparation, data acquisition, and isotopic labelling, etc^19,20^. First, current single-cell proteomic technologies use diverse sample preparation approaches (i.e., TEAB solution, DMSO solution, or water solution), data acquisition strategies (i.e., data-dependent acquisition or data-independent acquisition), and labelling strategies (i.e., label-free or isotopic labelling), resulting in severe batch effects^4,6,11^. Second, not all the constituting peptides’ contents are delivered to the mass spectrometry instrument due to the sample loss during sample preparation, which is a significant issue for single-cell proteomics. After the peptides are injected to the mass spec, their signal intensity would be influenced by their ionization efficiencies and peak selection of precursors for fragmentation in the DDA mode^19^, which will potentially lead to severe noise in single-cell proteomic data. Third, as for the bottom-up single-cell proteomics, the core quantification lies in the peptide level and a protein’s content is always obtained based on the simple average or median of its constituting peptides intensity detected in the mass spectrometry instrument. However, in practice, the current sequencing technologies are imperfect in quantifying peptide levels. It would be inaccurate to calculate the protein content only based on the average or median intensity of its peptides without considering the uncertainty of the measurement of the peptide intensities^8^ .Therefore, a more accurate way to construct the protein-level expression data is to assign uncertainty weights to peptides to compensate the measurement inaccuracy.

The existing single-cell proteomic data processing methods are mainly migrated from pipelines for scRNA-seq processing such as routine imputation (i.e., KNN imputation), batch correction (ComBat), and normalization^19,21,22^. However, those pipelines could not address the unique problems of data analysis on single-cell proteomic data^19^. First, KNN imputation could introduce big artifacts under severe batch effects in single-cell proteomic data. On the other hand, batch effect correction methods (ComBat) could not handle missing data without conducting imputation first. Hence, it would be problematic to conduct imputation and batch correction separately if the influence between them is ignored. Therefore, it is a significant challenge to consider the influence and interaction between these problems including data denoising simultaneously^19^. Second, current batch correction methods assume that there are the same sets of cells between two batches which might not be true when the cell types or cell states are unknown before the experiment. Third, single-cell proteomics exhibits a hierarchical structure that the proteins are constituted by peptides, and the protein contents are calculated based on the detected peptides’ signals. However, existing data analysis methods ignore the hierarchical information and the uncertainty of peptide quantification.

In this regard, we developed a novel deep graph contrastive learning framework for *s*ingle-*c*ell PROTeomics EmbeddINg (scPROTEIN) to tackle the uncertainty of peptide quantification, data missingness, batch effects and high noise in a unified framework by providing versatile cell embeddings. For the datasets provided with raw peptide signal intensities, we proposed a multi-task heteroscedastic regression model to estimate the uncertainty of peptide quantification and then aggregate the peptide content to the protein level in an uncertainty-guided manner. Then, we built a graph structure to characterize the single-cell proteomic data where the message-passing process considering co-expression patterns can help alleviate the data missingness problem and recover the missing values for low-abundance proteins. Graph contrastive learning model together with a designed alternated topology-attribute denoising module was developed, which can denoise the proteomic data and lead to an accurate representation. The discriminative property of contrastive learning and the well-denoised cell embedding can together alleviate the severe batch effect implicitly without knowing the prior knowledge of the dataset. Finally, the learned versatile cell embedding could be applied in various downstream tasks (i.e., cell clustering, batch correction and cell type annotation). We also performed experiments on antibody-based single-cell proteomic data to evaluate the effectiveness on clinical analysis. Moreover, scPROTEIN could be easily generalized to learn from spatial-resolved single-cell proteomic data.

## Results

### The overall architecture of versatile deep graph contrastive learning framework for single-cell proteomics embedding

The overall framework is illustrated in Fig. 1. For datasets provided with raw peptide-level profile, scPROTEIN starts from *stage 1* to learn the peptide uncertainty and obtain the protein-level expression profile in an uncertainty-guided manner. The protein-level data is then fed into *stage 2* and *stage 3* to learn the cell embedding by graph contrastive learning together with a data denoising module. The learned cell embedding can then be applied in a variety of downstream tasks.

**Figure 1.**
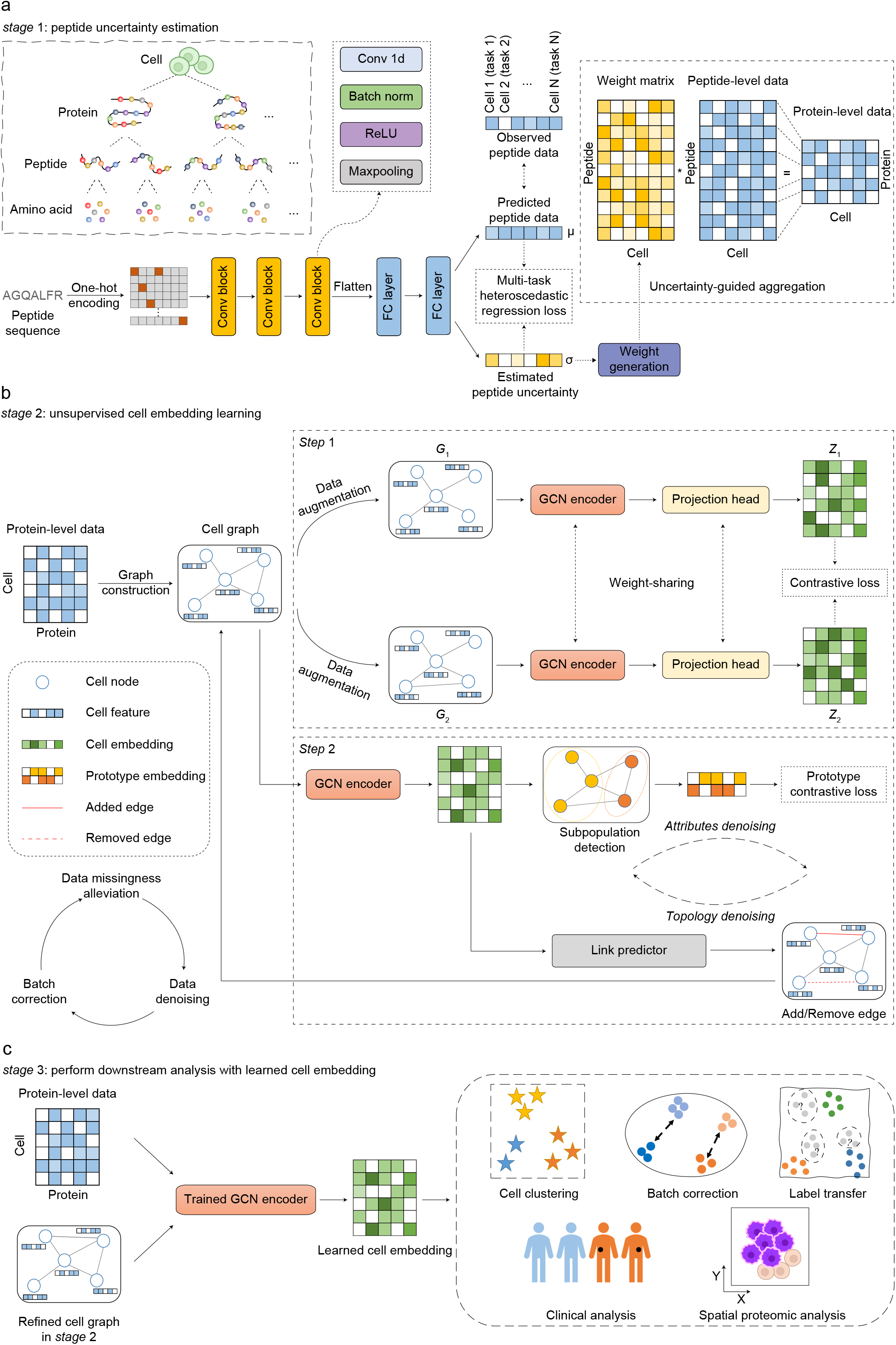
The architecture of our proposed method. **a**, The multi-task heteroscedastic regression model for peptide uncertainty estimation. The hierarchical structure for cells, proteins, peptides, and amino acids is shown in the left sub diagram. The uncertainty of peptide measurement is estimated in this stage and used to guide the aggregation of peptide levels for the corresponding protein. The network takes multiple peptide sequences as input in each iteration with a mini-batch training strategy. One peptide sequence is shown to illustrate the learning process. In the right panel, we show the aggregation process of using the learned uncertainty to get the protein-level content. **b**, The unsupervised cell embedding model based on deep contrastive learning. A cell graph is constructed using single-cell proteomic data where cells with similar protein content patterns are connected. Contrastive learning scheme is employed to generate cell embeddings. Attribute denoising and topology denoising are alternated to further improve the embedding. **c**, The inference of the pre-trained graph encoder to generate cell embedding, which is then applied to various downstream tasks.

The first stage is to aggregate the peptide-level intensities to determine the protein-level content guided by the uncertainty of peptide abundance quantification of mass spectrometry in single-cell proteomic data. We designed a multi-task heteroscedastic regression model to estimate the uncertainty of each peptide signal in each cell, which can partially reflect the quality of the signal (i.e., the amount of noise). Heteroscedastic regression model assumes that the uncertainty varies with different peptides. By considering the uncertainty estimation for all peptides in one cell as one specific task, our multi-task learning takes the estimation for all cells into consideration at the same time. The combination of heteroscedastic regression model and multi-task learning enables adaptively learning the uncertainty of each peptide across different cells. Based on the estimated uncertainty, we can then construct a weight for each peptide and perform uncertainty-guided aggregation over peptide-level data to obtain the protein-level data. By this way, peptide with higher quality can pose larger impact on the cell embedding and this process would benefit the model’s reliability and accuracy. Then we utilized the protein-level data to learn comprehensive cell embeddings in *stage 2*. For datasets provided directly with the reconstructed protein-level profile, scPROTEIN will start from *stage2*.

The topological structure of graph can help impute the missing data of single-cell proteomic data and improve the quality of learned embedding. Specifically, each node in the graph has specific semantic pattern which can be used to compensate each other during the graph learning process. To this end, in the second stage a cell graph is built where each node represents a cell and the initial node feature represents the proteomic data within the cell. There are three characteristics of our proposed graph: 1) in the contrastive learning framework, two different perturbed views are generated respectively and the mutual information of the same node between these two views is maximized, which can lead to robust representations to noise. 2) Besides, when dealing with data from different sources, the shared proteins are utilized to build a shared cell graph. Therefore, the batch effect between different sources can be implicitly alleviated by aligning the semantic information of the same cell type through contrastive loss. 3) Moreover, two denoising modules (attribute and topology) are alternated to mitigate the noise problem in the proteomic profile. During this *stage 2*, data missingness is alleviated, batch effect is implicitly corrected and the severe noise in proteomic data is largely removed.

In *stage 3*, the obtained topology-refined cell graph in *stage 2* and the single-cell protein-level data are fed into the trained GCN encoder in *stage 2*. Then the cell embedding is learned, which can be applied to various downstream tasks including cell clustering, batch correction, cell type annotation and clinical analysis. Moreover, our proposed methods can be easily extended to single-cell spatial proteomic data by constructing a cell graph based on the spatial cell proximity and learn a spatial informative embedding. A detailed description of the proposed method is in the Method Section.

To comprehensively evaluate the performance, we applied scPROTEIN in nine single-cell proteomic datasets (detailed in Supplementary Table 1). We first showed the overall learning workflow from *stage 1* to *stage 3* by taking a representative single-cell proteomic dataset SCoPE2_Specht^23^ as a concrete example. We qualitatively and quantitatively compared the cell clustering performance of scPROTEIN with other methods and showed the learned peptide uncertainty. Then we benchmarked five independent batch correction tasks across six single-cell datasets with different cutting-edge sequencing techniques (Scope2, pScope, plexDIA, N2, nanoPOTS, etc) and species (mouse and human cell lines). Besides, we investigated the application of scPROTEIN for single-cell clinical proteomic data in ECCITE-seq dataset^24^, which is an antibody-based single-cell proteomic dataset and collected from the patient with cutaneous T-cell lymphoma (CTCL). Furthermore, BaselTMA dataset^25^ which is consisted of spatially resolved proteomic data from breast cancer tumor biopsy slices was utilized to validate the scalability of scPROTEIN on single-cell spatial proteomics. Finally, a systematic analysis for hyperparameter sensitivity of our scPROTEIN model is presented in Extended Data Fig. 1 and Extended Data Fig. 2.

### Evaluation of cell clustering and the uncertainty of peptide quantification

We first evaluated the cell embedding learned by our proposed method on the cell clustering task and illustrated how scPROTEIN works from *stage 1* to *stage 3*. We applied scPROTEIN in SCoPE2_Specht dataset^23^, which quantifies 3042 proteins in 1490 cells via SCoPE2 sequencing technique. The existing single-cell proteomic data analysis pipeline^19^ employed KNN-based imputation and ComBat based batch correction (termed KNN-ComBat) for the routine data preprocessing. Beyond that, Markov affinity-based graph imputation of cells (MAGIC)^26^ is a manifold learning-based algorithm for denoising high-dimensional data which is commonly applied to single-cell RNA sequencing data. In this study, we used these two representative data processing methods as comparison method. From the clustering results in Fig. 2a-b, we can see that our graph-based embedding achieved the best performance on all the evaluation metrics on the SCoPE2_Specht dataset. Besides, we illustrated a concrete example of embedding learning process in Extended Data Fig. 3a.

**Figure 2.**
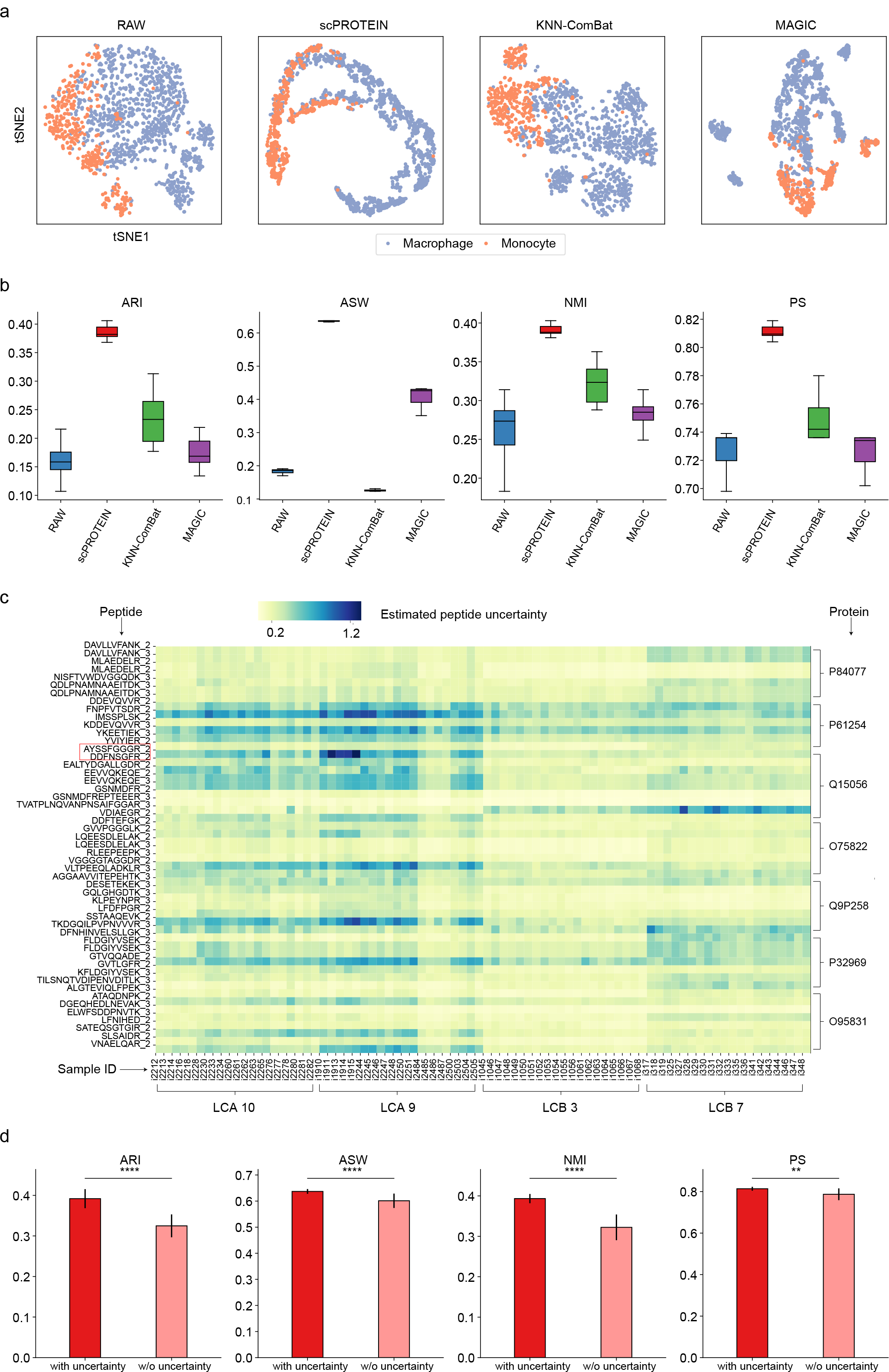
Cell clustering and peptide uncertainty estimation. **a**, The tSNE plot displaying the cell distribution with the embedding of our proposed method and comparison methods (KNN-ComBat and MAGIC) on SCoPE2_Specht dataset. The plots are colored by the cell types. RAW denotes the results of raw protein-level expression data. **b**, The ARI, ASW, NMI, and PS of our proposed method and comparison methods (KNN-ComBat and MAGIC) on cell clustering on SCoPE2_Specht dataset. RAW means the metric evaluated on the raw protein levels. The lower and upper hinges denote the first and third quartiles, with the whiskers in the range of 1.5-times the interquartile. **c**, The heatmap showing the estimated uncertainties of each peptide signal across cells, colored by the estimated uncertainty. The batch information and protein information are shown below the heatmap and on the right hand side of the heatmap, respectively. **d**, The ablation experiment on clustering performance based on our method with and without performing the uncertainty estimation of peptides in *stage 1* on SCoPE2_Specht dataset. Our method without uncertainty means that the protein content is obtained simply based on the median of its constituting peptides intensity. All data are given as means±sd. *, **, *** and **** indicate p-value<0.05, p-value<0.01, p-value<0.001 and p-value<0.0001 respectively in a two-sided Mann– Whitney test.

Furthermore, the entire framework is tailored for analysing the hierarchical data structure of single-cell proteomics where the expression of a protein can be aggregated by the levels of its digested peptides. The estimated peptide-level uncertainty reflects the data noise varied across samples and is visualized in Fig. 2c. As the digested peptides from different samples are pooled together after isotopic labelling and then detected in the mass spectrometry with consistent ionic behavior^4^, the peptides from the same run of experiments display similar uncertainty patterns. We can observe four distinct heatmap blocks along with the batch ID in Fig. 2c. While in the same proteins, the measurement uncertainties of different peptides vary due to their different behaviours in ionization, co-isolation, fragmentation, and sampler preparation loss^20,27^. For instance, “AYSSFGGGR_2” and “DDFNSGFR_2” are two constituting peptides of the same protein “Q15056” (in the red box), while they show quite different uncertainty patterns across batches and samples.

In addition, the ablation study in Fig. 2d displays the results of our method on SCoPE2_Specht dataset with and without uncertainty estimation in *stage 1*. It can be seen scPROTEIN performs significantly better after uncertainty adjustment. Besides, we compared the average signal to noise ratio within each cell type with other methods (Extended Data Fig. 3b), and the results prove the effectiveness of scPROTEIN in data denoising.

### Evaluation of batch correction with cell embeddings and label transfer

Due to the systematic biases in sample preparation, data acquisition, and labelling strategy, single-cell proteomic data faces more severe batch effects than scRNA-seq data. Therefore, we evaluated the robustness of our method to batch effect compared with existing methods on five independent experiments where the batch effects from different single-cell proteomic platforms hamper the clustering results of different cell types with the original protein level (Fig. 3a, c, Extended Data Fig. 4 a, Extended Data Fig. 5 a, c). We used different shapes to represent various cell types and the same colour in different shades to represent the same cell type from different sources. We first applied scPROTEIN to integrate two mouse cell datasets: N2^13^ (108 cells and 1068 proteins) and nanoPOTS^12^ (61 cells and 1225 proteins). The shared proteins (762 proteins) were used as the raw protein profile to build a shared cell graph. In Fig. 3a, the visualization of the raw data and processed data from comparison methods show a clear separation of the same cell type from both batches. After processed by scPROTEIN, cells within each cell type are pulled closer whilst those of different cell types remain properly separated. This indicates that scPROTEIN preserves the batch-invariant biodiversity. In addition, we show the comparison of raw protein data and the learned embedding by scPROTEIN in Extended Data Fig. 3c, which indicates that scPROTEIN can largely alleviate the batch effect.

**Figure 3.**
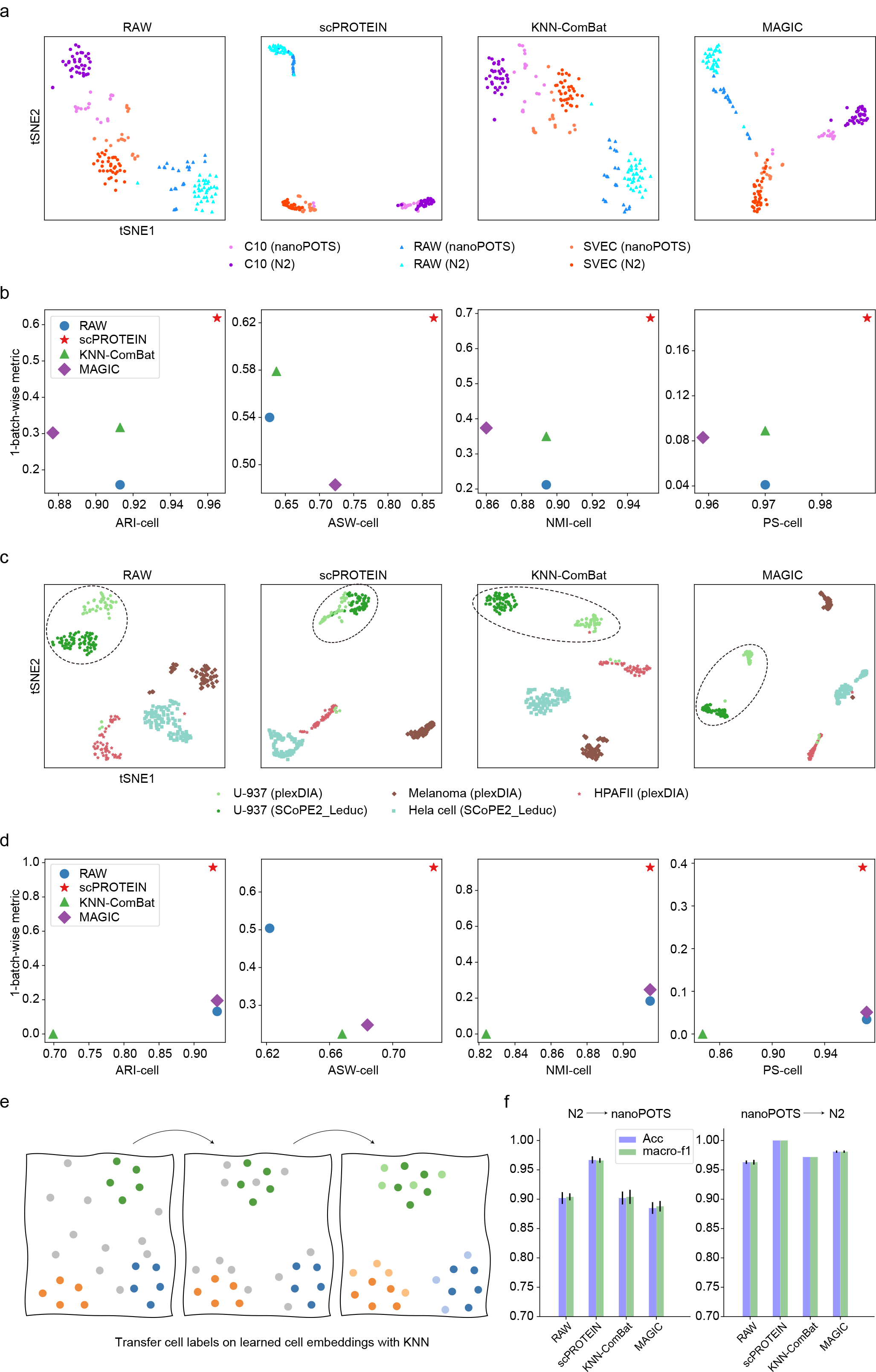
Batch effect correction and label transfer. **a**, The tSNE plot showing cells of N2 and nanoPOTS datasets, colored by data source and cell types. **b**, The ARI-cell, ASW-cell, NMI-cell, and PS-cell of our proposed method and comparison methods (RAW, KNN-ComBat, and MAGIC) with cell type labels as ground truth (x-axis) and 1-metrics of our proposed method and comparison methods (RAW, KNN-ComBat, and MAGIC) with batch labels as ground truth (y-axis) on N2 and nanoPOTs datasets. The result which is closest to the top right corner achieves the best performance. **c**, The tSNE plot showing cells of SCoPE2_Leduc and plexDIA datasets, colored by data source and cell types. U-937 is the shared cell type in the two datasets. **d**, The ARI-cell, ASW-cell, NMI-cell, and PS-cell of our proposed method and comparison methods (RAW, KNN-ComBat, and MAGIC) with cell type labels as ground truth (x-axis) and 1-metrics of our proposed method and comparison methods (RAW, KNN-ComBat, and MAGIC) with batch labels as ground truth (y-axis) on SCoPE2_Leduc and plexDIA datasets. **e**, The diagram showing the label transfer process based on the learned embedding. In the left panel, the gray dots represent cells with unknown labels from query set, and the other colored dots represent cells with known labels from reference set. When the batch effect is well removed (middle panel), the gray cells can then be annotated accurately by KNN (right panel). **f**, The histogram showing the accuracy and macro-f1 score of our proposed method and the comparison methods (RAW, KNN-ComBat, and MAGIC) on transferring labels from N2 to nanoPOTS (left panel) and from nanoPOTS to N2 (right panel). All data are given as means±sd.

For SCoPE2_Leduc^3^ (163 cells and 1647 proteins) and plexDIA^5^ (164 cells and 1242 proteins), the batch correction is more challenging since the two datasets have different sets of cell types. It can be seen from Fig. 3c that scPROTEIN still obtains a promising performance. Using scPROTEIN embedding, U-937 Cells from both sources were closely clustered while other cell types remained separable.

Besides, batch correction methods based on ComBat or other conventional methods may cause the problem of overcorrection for batch effects by removing the biodiversity of different cell types. To evaluate the performance of batch correction as well as the ability to preserve biodiversity within different cell types, we used cell-type-level metrics (ARI-cell, ASW-cell, NMI-cell, and PS-cell) to measure the cell type separation, and 1 - batch level metrics (1-ARI-batch, 1-ASW-batch, 1-NMI-batch, and 1-PS-batch) to measure the clustering of the same cell type between batches, similar to the metrics used in Tran et al^28^ and Li et al^29^. Results in Fig. 3b and Fig. 3d show the excellent balance of our method on correcting batch effect (1-batch level metrics) and maintaining the batch-invariant biodiversity of cells (cell type level metrics) compared to the comparison methods (Fig. 3b, d, Extended Data Fig. 4b and Extended Data Fig. 5b, d). Notably, the batch correction task for SCoPE2_Leduc and plexDIA is much more complex, as cells with the same cell type in both datasets are pulled closer due to batch removal, while the biodiversity of other different cell types have to be preserved and distinguished from noise. Hence, we employ Harmony^30^, the superior batch correction method for transcriptomics, to further evaluate the performance on this more difficult batch removal task tangled with the subtask of distinguishing noise from biodiversity. The cell clustering results of Harmony are ARI-cell = 0.682, ASW-cell = 0.424, NMI-cell = 0.722 and PS-cell = 0.823 and Harmony confuses different cell types from two datasets. In comparison, scPROTEIN has a better performance by drawing the same cell type closer while maintaining the biodiversity of other cell types under this tangled task.

We further performed label transfer to annotate cell types across N2 and nanoPOTs datasets which contain the same set of cell types. The label transfer was implemented by K- Nearest Neighbor (KNN) algorithm based on the batch correction results (Fig. 3e). The performance of label transfer is highly dependent on the effectiveness of batch corrections. Only when the batch effect is well corrected and the biodiversity is properly preserved, cells of unknown types can be moved closer to the correct cluster, thus enabling correct label transfer. As shown in Fig. 3f, when we take N2 as the reference set and nanoPOTS as the query set, the results indicate that the performance for cell type annotation of our method clearly outperforms other comparison methods on both accuracy (Acc=0.967) and macro-f1 score (macro-f1 score=0.966). Besides, our method can label all query cells correctly when transferring labels from nanoPOTS to N2 dataset (Acc=1.00, macro-f1=1.00).

### Application on clinical proteomic data

In this study, we briefly explored the application of our method on antibody-based single-cell proteomic data from clinical tissues in ECCITE-seq dataset^24^, which quantifies 49 marker proteins across 6500 cells from a healthy donor and 6500 cells from a patient with cutaneous T-cell lymphoma (CTCL). We visualized the UMAP of raw data and the learned representations in Fig. 4a and Fig. 4b, with cells from healthy donor in green and from cutaneous CTCL patient in red. The samples could be further grouped in 25 clusters with the embedding learned by scPROTEIN (Fig. 4c). The high expressed proteins of each cluster are shown in Fig. 4d. In addition, from Fig. 4e we can observe the proportion of cells from CTCL donor and from healthy donor varies in different subgroups. We selected three representative clusters and conducted detailed analysis for subgroups in Fig. 4f-n and more results can be found in Extended Data Fig. 6, in which pie plot indicates the proportion of CTCL samples and healthy samples (Fig. 4f, i, l), volcano plot depicts the differential expression proteins (Fig. 4g, j, m) and enrichment analysis indicates the gene ontology (GO) function for identified up-regulated proteins for CTCL cells (Fig. 4h, k, n). Based on the clustering results and corresponding subgroup analysis, we found several differential expressed proteins of the malignant CTCL cells^28^. For instance, CD62L (Fig. 4g) is the functional marker for circulating innate lymphoid cell precursors and correlates with the extent of lymphadenopathy, which could be applied as the diagnostic marker in inflammatory disorders^31,32^. For cluster 5 which overrepresents in CTCL sample, the volcano plot both indicates that CD62L is a potential functional and diagnostic marker for CTCL. As for cluster 6 which underrepresents in the CTCL sample, programmed death-1 (PD-1) is significantly upregulated in the cells from CTCL sample. The increased PD-1 level has been detected in various immune cells in CTCL patients and functioned in the attenuation of the immune response and anti-tumor immunity in the CTCL progression^33^. We also analysed the clusters which do not have significant variation between CTCL cells and control cells (i.e., cluster 2, Fig. 4l). CD2 is a fundamental gene that expresses on the surface of 80-90% of human peripheral blood lymphocytes, subtypes of NK cells and all mature T cells and those CD2+ cells within cluster 2 function as lymphocyte activation, immune system process and so on. Note that CD62L, the potential functional and diagnostic marker identified by our proposed method cannot be discovered by routine analysis employed on raw expression data in the original research on this dataset^24^.

**Figure 4.**
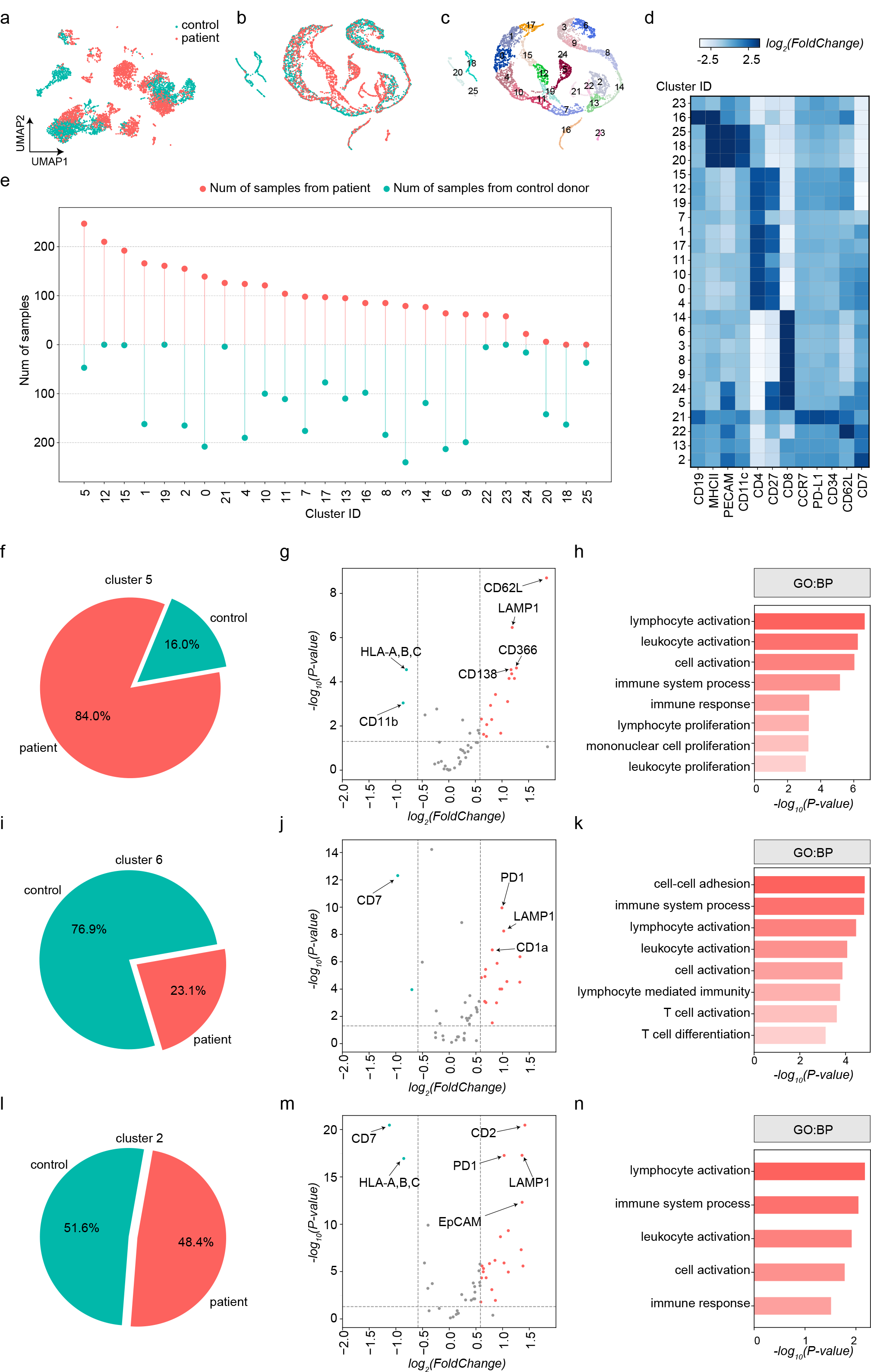
Application of learned embedding on clinical proteomic dataset. **a**, The UMAP for raw data showing the cells from healthy donor in green and cutaneous T-cell lymphoma (CTCL) patient in red. **b**, The UMAP for scPROTEIN showing the cells from healthy donor in green and cutaneous T-cell lymphoma (CTCL) patient in red. **c**, The UMAP for scPROTEIN showing the cells colored by clustering results. **d**, The heatmap showing the level of protein markers across clusters. **e**, The count numbers of cells from the control donor in green and CTCL donor in red for different clusters. **f**, The detailed ratio of cluster 5 cells in control donor and CTCL donor. **g**, The volcano plot showing the differential expression proteins found by contrasting healthy cells and CTCL cells in cluster 5. The dots in red and green represent the identified up-regulated and down-regulated proteins for CTCL cells, respectively. We used p-value = 0.05 and Fold Change = 1.5 as thresholds. **h**, Top GO terms in Biological Process (BP) for identified up-regulated proteins for CTCL cells in cluster 5. **i**, The detailed ratio of cluster 6 cells in control donor and CTCL donor. **j**, The volcano plot showing the differential expression proteins found by contrasting healthy cells and CTCL cells in cluster 6. **k**, Top GO terms in Biological Process (BP) for identified up-regulated proteins for CTCL cells in cluster 6. **l**, The detailed ratio of cluster 2 cells in control donor and CTCL donor. **m**, The volcano plot showing the differential expression proteins found by contrasting healthy cells and CTCL cells in cluster 2. **n**, Top GO terms in Biological Process (BP) for identified up-regulated proteins for CTCL cells in cluster 2.

### Application on spatial proteomic data

Spatial proteomic technologies are maturing and provide an increasing number of resources for the tumor microenvironment. We applied our method to single-cell resolved spatial proteomic datasets BaselTMA dataset^25^ by constructing a cell graph based on the spatial distance (Fig. 5a). As cells of the same cell types may locate in close proximity in the tissue, our method could enhance the cell embedding with the help of its spatial neighbor’s proteome. In this way, we can fully utilize both the spatial information and the protein profiling of the spatial proteomic data. The advantage of spatial proteomics is that it can reveal the spatial cell-cell interaction in the tumor microenvironment. Therefore, we analyze the tumor microenvironment with the learned embedding by scPROTEIN. Besides, the constructed spatial informative cell graph can be naturally used to quantify the spatial heterogeneity degree, which is utilized to estimate the spatial heterogeneity of the tumor microenvironment. In particular, a metric *shd* is defined to quantify the spatial heterogeneity degree and the detailed definition can be found in Evaluation Metrics in Methods. A high *shd* represents a highly compartmentalized phenotype and the tissue tends to be block-like. On the contrary, a low *shd* denotes a high level of spatial mixing.

**Figure 5.**
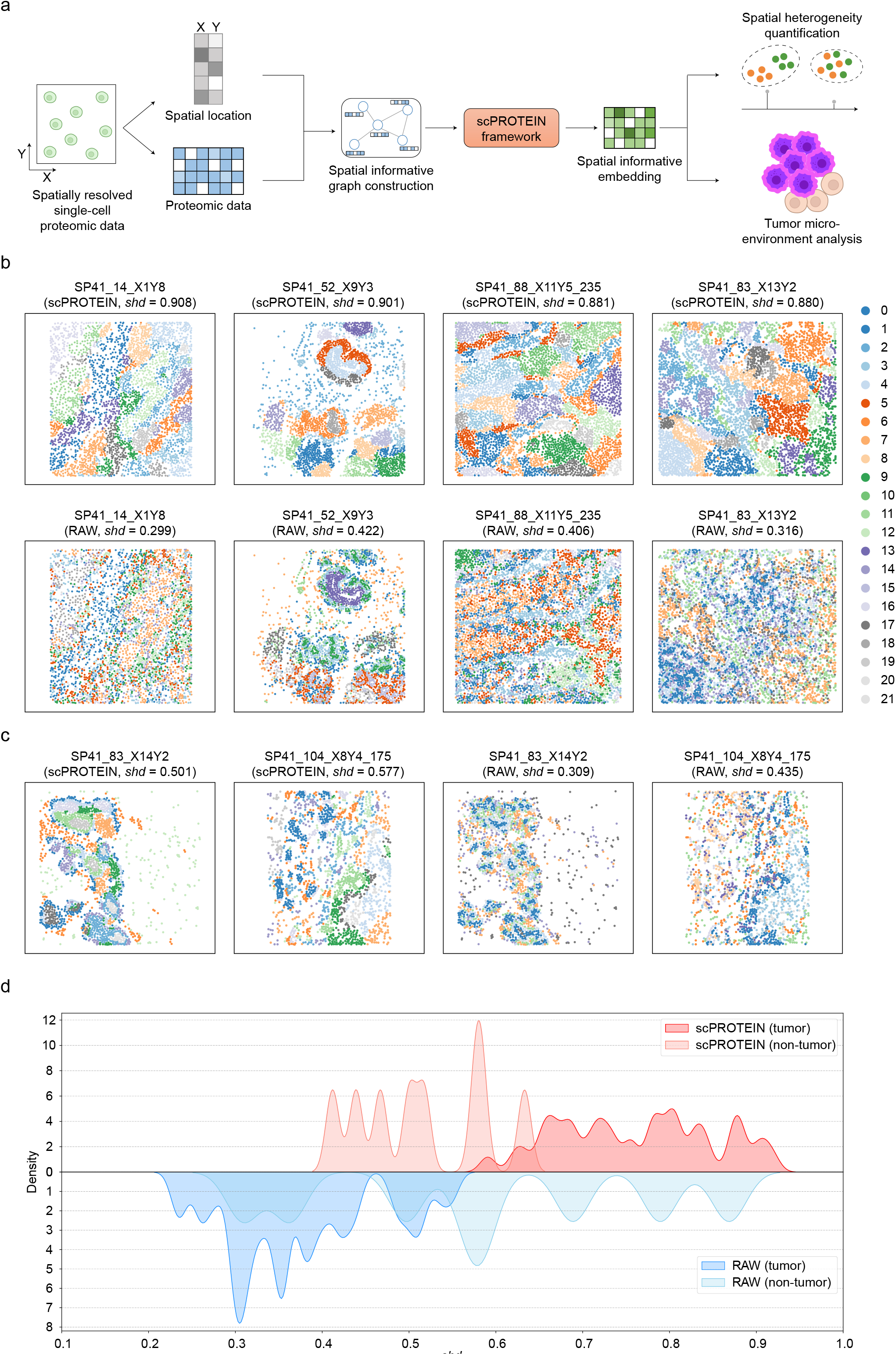
Application of our proposed method on spatial proteomic data. **a**, The diagram showing the application of our proposed method on spatial proteomic data. The spatial proteomic data is used to construct the spatial informative graph and then to infer spatial informative embedding for tumor microenvironment analysis and spatial heterogeneity degree quantification. **b**, The visualization of clusters on learned spatial informative embedding and the estimated spatial heterogeneity degree within tumor samples (top panel). Representative samples with high *shd* indicate a highly compartmentalized phenotype. The visualization of clusters on raw protein expression data and the estimated spatial heterogeneity degree within tumor samples (bottom panel). *shd* is calculated based on the graph built in scPROTEIN. **c**, The visualization of learned spatial informative embedding and the estimated spatial heterogeneity degree within non-tumor samples (left panel). Representative samples with low *shd* indicate a high level of mixing between different cells. The visualization of clusters on raw protein expression data and the estimated spatial heterogeneity degree within non-tumor samples (right panel). *shd* is calculated based on the graph built in scPROTEIN. **d**, The density plot of *shd* for scPROTEIN and raw expression data. The *shd* of scPROTEIN for tumor and non-tumor slices show different ranges, while in raw expression data the ranges are overlapped.

Here the cell clusters identified by scPROTEIN from tumor slices shown in Fig. 5b (top panel) indicate the highly compartmentalized phenotype and exhibit high *shd* scores (range from 0.880 to 0.908). In contrast, the spatial proteomic data from the normal slices retains relatively low *shd* scores (0.501 and 0.577), which indicates the high levels of inter-cell-type mixing (Fig. 5c, left panel). The *shd* scores indicate two types of samples showing different spatial heterogeneity signatures (p-value = 0.0015, two-sided Mann–Whitney test). The observation is consistent with the previous clinical study on breast tumor^25,34^.

Besides, in Fig. 5b (bottom panel) and Fig. 5c (right panel), we depict the results by directly clustering on raw single-cell protein data for tumor slices and normal slices, respectively. It can be seen that for two different types of slices, the results do not show any distinct signatures. We further show the density plot of *shd* scores across different slices for both scPROTEIN and raw expression data in Fig. 5d. For scPROTEIN the *shd* for tumor slices and non-tumor slices show different ranges, while in raw data the ranges for both types are overlapped. This confirms the effectiveness of scPROTEIN to discriminate the compartmentalized and highly mixed spatial phenotypes in spatial resolved single-cell proteomic data. More results are shown in Extended Data Fig. 7 a, b.

## Discussion

Although the widely used CITE-seq technology facilitates the detection of cell surface proteins at scale, it suffers from the limitation of detectable protein numbers and types. The mass-spec based proteomics provides complementary advantages in detectable protein numbers and types (1000-3000 intracellular proteins) which are critical for analysing cellular function and disease progression. However, at the same time, it is a significant challenge to simultaneously address the peptide uncertainty estimation, data missingness, batch effect and data noise problem in mass-spec based single-cell proteomic data. Most of the existing methods only address one or two problems, but generally speaking, because these problems are often tangled each other, previous methods are not effective to be used alone or simply combined.

To this end, we presented a versatile deep graph contrastive learning model scPROTEIN that is tailored for single-cell proteomics embedding and solved the whole set of problems in a unified framework. First, with the uncertainty of peptide signals estimated by the multi-task heteroscedastic regression model, our method could aggregate peptide-level content to protein content. In this way, we could fully utilize and explore the available hierarchical information of single-cell proteomic data. The nascent design of uncertainty estimation in peptide intensity exactly meets the urgent need of the single-cell proteomics community^8^. Then, the cell graph constructed by cellular expression of proteomic displayed the cell’s similarity and the data missingness problem could be implicitly alleviated by propagating information from neighbour cells. scPROTEIN was trained based on contrastive learning, which can lead to a perturbation-resistant cell embedding. To further obtain a more robust representation, we designed an alternated topology-attribute denoising module. The noisy raw expression profile could be largely cleaned, and the initial constructed cell graph topology was enhanced. Moreover, contrastive learning is capable of pull the cells with similar pattern closer and push quite different cells apart, which yields a discriminative ability for batch correction and promotes data completion as well as denoising. To the best of our knowledge, it is the first time to establish the cell embeddings of single-cell proteomic data to address all above problems, providing a novel insight into exploring the embedding of single-cell proteomic data in a data-driven manner.

Extensive experiments proved the versatile application and superior performance of the embeddings learned by our method on both mass-spec-based and antibody-based proteomics. Benefiting from the compact embedding, scPROTEIN demonstrates a promising performance compared with the existing single-cell proteomic data processing pipeline and other comparison methods in cell clustering, batch correction and cell type annotation on a large number of single-cell proteomic datasets. Moreover, scPROTEIN exhibits wide applicability such as clinical analysis and single-cell spatial resolved proteomics data analysis. The application of our method on clinical single-cell proteomic data would be further explored as the exploration of the single-cell proteomics application in clinical practice. In terms of the generalization capability, scPROTEIN could be easily extended to spatial-resolved single-cell proteomic data. Building cell graph based on cell proximity yields a spatial-informative embedding and the constructed graph could be naturally used to quantify the spatial heterogeneity degree. It could be foreseen that with the rapid development and application of single-cell proteomic technologies, our method would play an increasingly important role in various scenarios of single-cell proteomic data analysis.

Despite the above advantages, there still exist some limitations in our investigation. First, some sequencing platforms provided raw profile data directly from the protein-level, in which *stage 1* tailored for the hierarchical proteome structure cannot be employed. Second, since there is no ground truth for peptide uncertainty, the performance of peptide uncertainty estimation is evaluated indirectly by downstream tasks. Third, the existing pipelines for single-cell proteomic data are scarce, and therefore few baseline methods can be used for comparison. In addition, with the rapid development of graph model, there will emerge more informative graph construction manners in the future, which could further enrich our scPROTEIN model.

## Methods

### The proposed method

#### Estimation of the uncertainty of the peptide-level intensity

In the previous study^4,23^, the detection of peptide sequence is considered to be deterministic. However, it has been shown there is a probability that the detected signal is not accurate^8^. Therefore, it is necessary to estimate uncertainties of peptide-level signal and obtain a more informative protein-level data.

To this end, we develop an uncertainty-aware framework in scPROTEIN *stage 1* to provide a measure for peptide uncertainty estimation via multi-task heteroscedastic regression motivated by previous researches^35^. As shown in Fig. 1a, the model takes the original peptide sequences as input and outputs the estimated uncertainty. We employ Convolutional Neural Network (CNN) as the backbone framework.

Specifically, for each certain detected peptide sequence, we first use one-hot encoding to generate a matrix of size 20 × *peptide*_*length*, where 20 is the total number of common amino acid types and *peptide*_*length* is the length of the input sequence. Besides, since the sequence lengths are variable, we perform a padding operation to align all the one-hot matrices to the maximum sequence length, which facilitates the subsequent encoding process.

The one-hot encoded matrices then go through three convolutional blocks as shown in Fig. 1a, where each block consists of one-dimensional convolution layer, batch normalization layer, Rectified Linear Unit (ReLU) activation function and max-pooling layer. The outputs are then flattened and pass through two fully connected layers. Finally, we design two independent dense layers to obtain the predicted expression vector *μ* and estimated uncertainty *σ* respectively, and the loss function for training the whole peptide uncertainty estimation framework is:

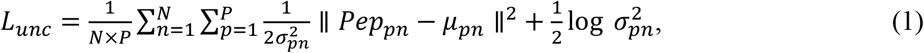

where *μ_pn_* and *Pep_pn_* represent the predicted and observed expression value of peptide sequence *p* in cell *n*, respectively. *σ_pn_* represents the uncertainty for peptide sequence *p* in cell *n*. It can be seen that this loss function is composed of two terms and each term involves *σ_pn_*, which measures the data uncertainty and also plays the role as a balance factor. Specifically, to reduce the loss, the model cannot output an overly small *σ_pn_* value since the term 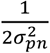 will explode. On the other hand, the model cannot predict an extremely large *σ_pn_* either since the second term will increase. Hence, the model will output a reasonable uncertainty estimation to balance both terms in the loss function. Besides, ∥ *y_pn_* − *μ_pn_* ∥^2^ term is the Mean Square Error (MSE) loss between the predicted and the observed expression values. Only when this term is small enough, scPROTEIN can offer a small uncertainty prediction *σ_pn_*, which indicates that the model is confident about the current prediction of peptide expression value.

#### Protein-level expression matrix aggregated from peptide estimation

Once the peptide uncertainty is estimated, we can compute the protein-level data in an uncertainty-guided manner as shown in the right side of Fig. 1a. A peptide signal with higher uncertainty estimation value indicates a noisy signal and should be assigned a smaller weight. In contrast, a peptide signal with lower uncertainty tends to be a highly confident signal and should be given a larger weight. Therefore, we can obtain the protein-level expression matrix *X* with its each entry as:

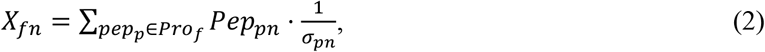

where *X_fn_* represents the aggregated expression value of protein *f* in cell *n* and *pep_p_* ∈ *Pro_f_* denotes peptide sequence *p* from protein *f*. The obtained expression protein-level expression matrix is then fed into the *stage 2* of the scPROTEIN framework as the initial feature matrix.

#### Graph construction

To make full use of the single-cell proteomic data, we convert the expression matrix into an undirected and unweighted cell-cell graph *G* = (*V*, *E*, *X*). *V* = {*v*_1_, *v*_2_, …, *v_N_*} is the set of cell nodes and *N* denotes the number of cells. *E* = {*e_ij_*} ⊆ *V* × *V* represents the set of all edges. We take the protein-level expression data as the feature matrix *X* = {*x*_1_, *x*_2_,…, *x_N_*} ∈ ℝ^*N*×*F*^, and *F* denotes the dimension of feature vectors which is also the number of proteins.

To obtain the cell graph topology, we first calculate the cell-wise similarity matrix *S* by Pearson correlation coefficient (PCC) based on the expression feature vectors. *S* is formally stated as:

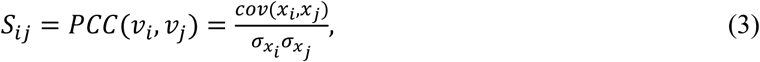

where *cov* and *σ* denote the covariance and the standard deviation respectively. Then we set a threshold *h* to construct the cell graph and the topological structure of cell graph *G* can be specified by a symmetric adjacency matrix *A* ∈ ℝ^*N*×*N*^:

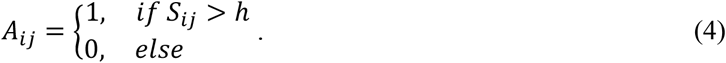

The obtained binary adjacency matrix is then used to perform the following graph learning. Besides, our scPROTEIN method can also be easily extended to single-cell spatial proteomics. With the provided spatial location information, scPROTEIN can construct cell graph based on spatial proximity. Specifically, the similarity matrix *S* now will be calculated based on the Euclidean distance computed from the spatial coordinates as:

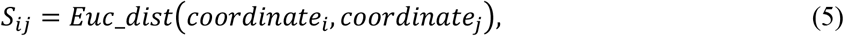

and we can similarly obtain the spatial informative cell graph topology by setting the threshold as mentioned above but *A_ij_* = 1 when *S_ij_* < *h*. Noteworthy, our constructed spatial informative cell graph can subsequently be used to quantify the spatial heterogeneity degree.

As the protein profile data are noisy, the constructed cell graph cannot be completely accurate. Therefore, we apply two data augmentation strategies and a special denoising module to solve this problem and learn robust embeddings.

#### Graph contrastive learning for accurate cell embedding

##### Model architecture

Since label information is always scarce and task-specific, we propose to learn embeddings in an unsupervised manner for better model generalizability. Motivated by the previous study^36^, we propose a deep graph contrastive learning framework to learn the comprehensive low-dimensional cell embeddings, which receives the cell graph *G* and expression feature matrix *X* as input. Specially, in order to deal with the noisy single-cell proteomic data caused by limitations of sequencing technology, we design a novel alternated topology-attribute denoising module, which yields a more informative and noise-resistant cell embedding. Overall, scPROTEIN contains four major components as shown in Fig. 1b: 1) Data augmentation, 2) GCN-based graph encoder, 3) Node-level graph contrastive learning and 4) Alternated topology-attribute denoising module. Next, we will delve into the details.

##### Data augmentation

Contrastive learning aims to learn the invariant representation between similar and dissimilar data pairs. To produce the similar pairs, we employ the data augmentation to generate different views of the data. Here we adopt two types of graph augmentation techniques: drop-edge^37^ and mask-feature.

For drop-edge, we randomly remove existing edges from *E* with a given ratio *p_de_*. In particular, we sample an indicator matrix *R* ∈ ℝ^*N*×*N*^ to decide which edges will be removed. *R_ij_* can be expressed as:

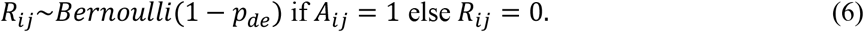

The perturbed adjacency matrix can be obtained by:

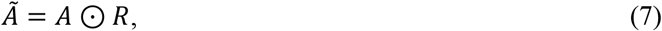

where ⊙ represents the Hadamard product operator.

For mask-feature, an indicator vector *M* ∈ ℝ^*N*×1^ is generated with each entry *M_i_*~*Bernoulli*(1 − *p_mf_*), where *p_mf_* is a given mask-feature probability. The masked feature matrix is expressed as:

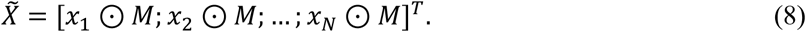

[∙ ; ∙] represents the vector concatenation operator. In each training iteration, we employ the data augmentation to generate two different but correlated graph views 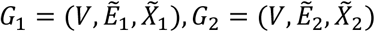 as shown in Fig. 1b *step 1*.

##### GCN-based graph encoder

After the augmented graph views *G*_1_ and *G*_2_ are prepared, we learn the cell embedding based on node-level graph contrastive learning. Since Graph Convolutional Network (GCN)^38^ provides a powerful learning model and is able to extract comprehensive embeddings for graph-structured data, we adopt GCN as the feature extractor to learn the latent pattern for cell nodes. Concretely, GCN performs convolution operation on the graph and iteratively updates the node representations by message passing among the neighbourhoods. Taking the graph topology structure and the feature matrix as input, the low-dimensional node representation can be learned by a *L*-layer GCN as:

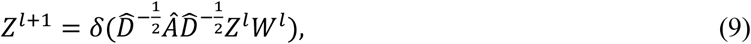

where *Â* = *A* + *I* and *I* is the identity matrix. 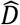 is the degree matrix of *Â* with 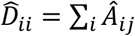. *W^l^* is the layer-specific trainable weight parameter and *Z^l^* represents the embedding matrix obtained by layer *l*. We use PReLU (Parametric Rectified Linear Unit) as the non-linear activation function *δ*(∙). For two augmented graph views, we adopt a weight-sharing GCN encoder to generate the embedding matrix *Z*_1_ and *Z*_2_ for each view respectively. We set the dimension of all GCN layers *d* and the dimension of output embedding matrix is ℝ^*N*×*d*^.

##### Node-level graph contrastive learning

The obtained output *Z*_1_ and *Z*_2_ are then fed into a weight-sharing projection head *g*, which is used to project the embeddings from two views into a common latent feature space where the contrastive loss is constructed. *g* is implemented by a two-layer Multiple Layer Perceptron (MLP). By calculating the contrastive loss in the projected latent space, scPROTEIN can obtain a better representation ability^39,40^.

Then we try to maximize the agreement of the representations between two generated views of the same node. Taking node *i* as an example, its embedding *z*_1*i*_ from view *G*_1_ can be regarded as an anchor and embedding *z*_2*i*_ from another view *G*_2_ will be treated as a positive sample. Naturally, the embeddings of other cell nodes will be taken as negative samples. We want to maximize the agreement between positive pairs (anchor and its positive sample) and minimize the agreement between negative pairs (anchor and its negative samples). Here we adopt the cosine similarity function cos (∙) to measure the similarity between samples and we can arrive at the modified infoNCE loss^41^ for positive pair (*z*_1*i*_, *z*_2*i*_) as:

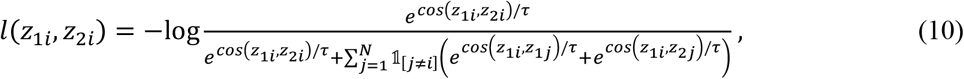

where *τ* is the temperature parameter and 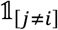 denotes the indicator function. (*z*_1*i*_, *z*_1*j*_) is the negative pair from the same augmented view and (*z*_1*i*_, *z*_2*j*_) is the negative pair from different views. Since the two generated views can be switched, we can finally obtain the overall contrastive loss expressed as:

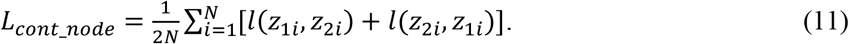

By minimizing this node-level contrastive loss, positive node pairs are drawn closer and negative node pairs are pushed apart. The biodiversity of each cell node can be well preserved and the batch effect can be implicitly corrected, which results in a more robust embedding.

##### Alternated Topology-attribute denoising module

Compared to single-cell transcriptomic data, proteomic data has a more severe noise problem due to its specific sample preparation process. Therefore, in order to obtain a noise-resistant cell embedding, we design an alternated attribute-topology denoising module, which can be decoupled into the following two alternated major steps.

###### Attribute denoising

To alleviate the severe noise problem in the single-cell proteomic profile, we develop an attribute denoising module based on prototype contrastive learning. As investigated in the previous study^42^, the prototypes^30^ which are far away from the cluster boundaries are not vulnerable to noise compared to those near the boundary. Based on this observation, we propose to exploit the away-from-boundary prototypes to update other noisy samples.

As shown in Fig. 1b *step2*, at each training epoch, we perform subpopulation detection via K-means algorithm^43^ based on the current learned embeddings with a pre-defined cluster number *K*. After K-means algorithm converges, we can assign each node with its cluster label *cluster_i_*. Then we naturally take *K* cluster centers {*c*_1_, *c*_2_, …, *c_K_*} as the prototype representation since these centers are usually far away from the cluster boundary and more confident about their cluster label. Formally, cluster center *c_k_* is calculated by the average of samples with cluster label *k*:

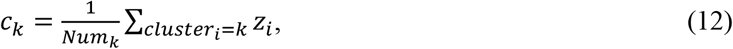

where *Num_k_* denotes the total number of samples with cluster label *k*. *z_i_* is the learned cell embedding for node *i* without data augmentation. This noise-resistant knowledge is then exploited to update the whole node representation matrix by the designed prototype contrastive loss as:

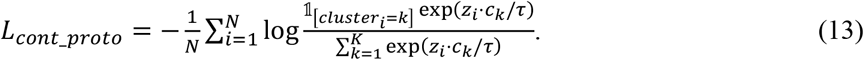

By minimizing this objective function, the similarity between each node and its cluster center will be enlarged. Each prototype is used to enhance the information of its surrounding samples. Finally, we can obtain a denoised cell representation after an iterative training process. Besides, we further confirm that our scPROTEIN model is robust to the choice of the cluster number *K* in Extended Data Fig. 2.

###### Topology denoising

To fix the existing inaccurate edges and further enhance the graph topology, we develop a topology denoising module. The graph learning process is highly dependent on the graph structure. Hence the noise exists in the graph topology can seriously degrade the model performance. In the initial cell graph, there may be some vital edges that have not been mined. On the other hand, some established edges may be pseudo edges which may bring in noisy information. With the learning of the network and the process of attribute denoising, the characteristics of the nodes are continuously enhanced and cleaned, and the semantic information of each cell node becomes clearer. Therefore, the edges in the network should also be alternately updated accordingly.

To this end, we design a link predictor to alternately denoise the topology structure, and in turn improve the quality of the learned embeddings. Considering the embedding matrix in iteration *t* as *Z*^(*t*)^, we can obtain the corresponding similarity matrix *S_ij_*(*t*) based on pair-wise Pearson correlation coefficient. Then we choose *M* edges with the highest probability in *S_ij_*(*t*) as the set *Edge*_*add*^(*t*)^, and *M* edges with the lowest probability score as the set *Edge*_*remove*^(*t*)^. The current adjacency matrix *A_ij_*(*t*) can be refined based on these two sets. We suggest that high probability edges should be added to enhance the graph representation ability, while low probability edges should be removed to eliminate the topological noise. Thus, we can obtain the updated adjacency matrix in iteration *t* as:

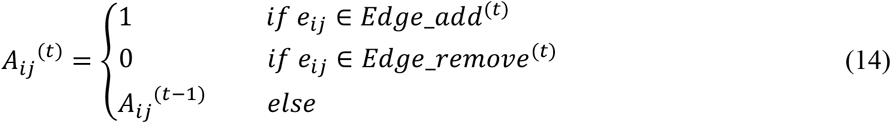

As shown in Fig. 1b *step 2*, the updated adjacency matrix will pass through the next training iteration for data augmentation and attribute denoising. Then the denoised node representation will in turn be used to update the topological structure. Through such an alternated update mechanism, we largely relieve the impacts of noise problem in single-cell proteomic data and derive a more robust cell representation.

##### Overall loss function

The overall loss function of scPROTEIN model consists of two parts, which is expressed as:

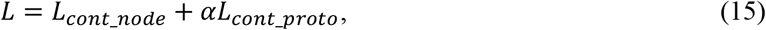

where *α* is a balance factor which is 0.05 used in the experiments. We minimize this loss function to train the model parameters. Adam^44^ is used as the optimizer with a learning rate 0.001.

#### Generation of cell embedding

After the model achieves convergence, we can obtain the trained GCN encoder and the refined cell graph. As shown in Fig. 1c, the learned cell embedding *Z* is generated on the refined graph topology in *stage 2* with the informative trained GCN encoder. The obtained embedding has largely mitigated this set of tangled problems, including the data missingness, batch effects and noise problem.

Therefore, it can be used in a variety of applications, such as cell clustering, batch correction, label transfer, clinical analysis and spatial analysis. It is worth noting that data augmentation is only used during the training process. During the inference process, none of the data augmentation techniques is exploited.

### Datasets

#### The single-cell proteomic datasets

##### SCoPE2_Specht^23^

SCoPE2_Specht is a representative single-cell proteomic dataset which quantifies 3042 proteins in 1490 cells via SCoPE2 sequencing technique. It contains two cell types: Monocyte and Macrophage cells. Noteworthy, since Monocyte cells can be differentiated into Macrophage-like cells when polarizing cytokines are absent, the characteristics of these two cell types can be quite similar and result in a more difficult cell clustering task. We downloaded the data from https://scp.slavovlab.net/Specht_et_al_2019.

##### nanoPOTS^12^

nanoPOTS dataset was prepared by nanoPOTS technology (nanodroplet processing in one-pot for trace samples). It quantifies 1225 proteins in 61 cells which are composed of C10 cell, RAW cell and SVEC cell. Data was downloaded at MassIVE data repository with ID: MSV000084110.

##### N2^13^

N2 dataset sampled from mouse blood contains 108 cells and 1068 proteins. In N2 dataset, there are three cell types (C10 cell, RAW cell and SVEC cell) as well. We downloaded the data from MassIVE data repository with ID: MSV000086809.

##### SCoPE2_Leduc^3^

The SCoPE2_Leduc dataset was downloaded from https://scp.slavovlab.net/Leduc_et_al_2021. It contains 163 cells over 1647 proteins which is generated by SCoPE2. In SCoPE2_Leduc dataset, there are two cell types: Hela cell and U-937 cell.

##### plexDIA^5^

We downloaded plexDIA dataset from https://scp.slavovlab.net/Derks_et_al_2022. It contains three cell types, which are Melanoma cell, U-937 cell, PDAC (HPAFII) cell.

##### pSCoPE_Huffman^6^

We obtained the pSCoPE_Huffman dataset from https://scp.slavovlab.net/Huffman_et_al_2022. This dataset is sampled by pSCoPE and consists of 163 cells and 1647 proteins. It contains three cell lines of PDAC cell: CFPACI, HPAFII and BxPC3.

##### pSCoPE_Leduc^45^

pSCoPE_Leduc dataset was downloaded from https://scp.slavovlab.net/Leduc_et_al_2022. It is generated by pSCoPE technique and 1543 cells over 2844 different proteins are quantified. Melanoma cell and U-937 cell are two cell types of pSCoPE_Leduc dataset.

#### The single-cell clinical proteomics datasets

##### ECCITE-seq^24^

The ECCITE-seq dataset was downloaded from Gene Expression Omnibus (accession number: GSE126310). It contains 49 surface protein markers which is detected by cellular indexing of transcriptomes and epitopes by sequencing (CITE-seq). The human peripheral blood mononuclear cells (PBMC) in ECCITE-seq are from a healthy control donor or a patient with cutaneous T-cell lymphoma (CTCL). After removing cell doublets, there remain 2767 cells from the control donor and 2634 cells from the CTCL donor which are then used for exploration of downstream clinical analysis based on single-cell proteomic data.

#### The spatial proteomics datasets

##### BaselTMA^25^

The BaselTMA dataset was downloaded at Zenodo (https://doi.org/10.5281/zenodo.3518284) and includes 281 patients with breast cancer. The imaging mass cytometry technique was used to quantify 38 marker proteins and spatial tissue image at the same time. This results in that each TMA is equipped with spatially resolved single-cell location information together with the protein quantification matrix. Each tissue slice contains a 0.8-mm tumor core or a corresponding healthy breast tissue.

### Data preprocessing

The raw single-cell proteomic data is firstly normalized to the library size and then log2 transformed with pseudo count 1. This process is implemented via SCANPY (https://pypi.org/project/scanpy/) package. Besides, for ECcite_seq dataset, we remove the cell doublets as the authors suggested by a python package scrublet^46^ (https://github.com/swolock/scrublet).

### Baseline methods

#### KNN-ComBat^19^

The imputation step with KNN is implemented by python package sklearn (https://scikit-learn.org/stable/) and ComBat for batch correction is implemented by scanpy.pp.combat module via SCANPY (https://pypi.org/project/scanpy/) package.

#### MAGIC^26^

MAGIC is downloaded from https://github.com/KrishnaswamyLab/MAGIC.

### Evaluation experiments

#### Performance on the cell clustering

We conducted cell clustering experiment on SCoPE2_Specht, which is a representative single-cell proteomic dataset. We first exploited scPROTEIN *stage 1* to estimate the uncertainty of the peptide signal. The obtained uncertainty estimation matrix together with the original peptide-level expression data were then combined to produce the protein-level expression data, which was then fed into scPROTEIN *stage 2* and *stage 3* to learn the informative cell embeddings. The embedding dimension was reduced by using top 50 principle components selected by PCA (Principal Component Analysis) and we then used tSNE (t-distributed stochastic neighbor embedding) algorithm to further reduce the dimension to 2 for 2-dimensional visualization similar to Li et al^29^. The PCA and tSNE function were both implemented via python package sklearn.manifold. Furthermore, to assess how well scPROTEIN performs by the aid of the uncertainty-guided aggregation for protein-level data, we conducted an ablation study to compare the performance of scPROTEIN with and without *stage 1* (Fig. 2d).

#### Performance on the batch correction

To evaluate the ability of our proposed methods on cross-cohort data with batch effects from various single-cell proteomic sequencing techniques and platforms, we conducted five different experiments in two levels. In the first level, we performed experiments on N2 and nanoPOTS datasets with exactly the same set of cell types (Fig. 3a, b). In these two datasets, both have three cell types: C10 cell, RAW cell, and SVEC cell. In the second level, we test the model’s ability to reconcile the same cell type (cell line) from different sources while maintaining the biodiversity of different cell types.

Specifically, we corrected the batch effect of SCoPE2_Leduc and plexDIA in Fig. 3c, d (merging U-937 cell), pSCoPE_Huffman and plexDIA in Extended Data Fig. 3a, b (merging PDAC (HPAFII) cell), pSCoPE_Leduc & plexDIA in Extended Data Fig. 4a, b (merging Melanoma cell and U-937 cell), and pSCoPE_Leduc & SCoPE2_Leduc in Extended Data Fig. 4 c, d (merging U-937 cell). In each batch correction experiment and for all comparison methods, the shared proteins were exploited for analysis. We reported the mean values of each metric across 10 runs in each experiment.

#### Performance on the cell type annotation

The cell type annotation (label transfer) task was conducted on N2 and nanoPOTS dataset. After correcting the batch effect via scPROTEIN, the cell labels from one dataset can be naturally transferred to another (Fig. 3e, f). KNN algorithm was employed to inference the labels of query set from reference set in the learned latent space.

### Application

#### Performance on clinical single-cell proteomic data

Samples from ECCITE-seq data were clustered based on the learned embeddings by Leiden algorithm (resolution=0.6), which was implemented via scanpy.tl.leiden. Samples were clustered into 25 subgroups. We can observe the different sample proportions and distinct protein signatures in various clusters using the cell embedding. The Differential expression protein analysis was carried out using rank_gene_groups function in SCANPY package with the t-test statistical method. The Functional profiling of Gene Ontology (GO) for each subpopulation was performed by the gProfiler^47^ (https://biit.cs.ut.ee/gprofiler/gost).

#### Performance on spatial single-cell proteomic data

BaselTMA dataset was exploited to confirm the scalability for scPROTEIN on single-cell spatial proteomics. In each TMA slice, we constructed the cell graph, where node attributes are the expression profile and edges between cells are established based on spatial cell location. Then the spatial informative embedding can be naturally obtained via scPROTEIN model. The learned embedding was then clustered via Leiden algorithm (resolution=0.6).

### Evaluation Metrics

ARI^48^ (Adjusted Rand Index), ASW^49^ (Average Silhouette Width), NMI^50^ (Normalized Mutual Information) and PS^51^ (Purity Score) were used for the evaluation of cell clustering and 1 - batch wise metrics (1-ASW, 1-ARI, 1-NMI, and 1-PS) were used to measure the performance on batch correction of same cell type. Similar to ^28,29^, we applied these metrics on the transformed data by tSNE. Accuracy and macro-f1 score were applied in cell type annotation experiments. Average signal to noise ratio was exploited to evaluate the data quality and quantify the data noise within each cell type. For all metrics, higher score represents better performance. Metrics are defined as follows:

#### ARI^48^

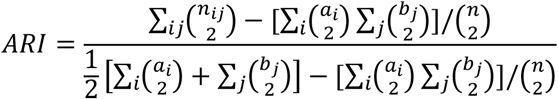

where n_ij_ denotes the number of cells which are assigned to cluster i and j based on the true labels and the clustering labels respectively. a_i_ is the number of cells from cluster i based on the true labels and b_j_ is the number of cells assigned to cluster j according to the clustering labels.

#### ASW^49^

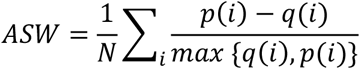

*q*(*i*) represents the average distance of cell *i* to other cells in the same cluster and *p*(*i*) is the average distance between cell *i* and other cells in the nearest cluster. *N* is the total number of cells.

#### NMI^50^

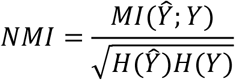

*MI*(*Ŷ*; *Y*) denotes the mutual entropy between the predicted categorical distributions *Ŷ* and the true clustering categorical distributions *Y*. *H* is the Shannon entropy.

#### PS^51^

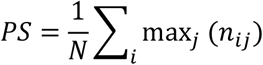

*n_ij_* denotes the number of cells assigned to cluster *i* whose ground-truth label belongs to partition *j*. The purity score quantifies the extent to which a cluster contains cells from only one partition.

#### Acc

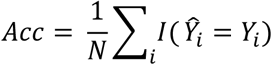

*I*(∙) denotes the indicator function.

#### macro-f1

The macro-f1 score was calculated by the function f1_score with average=“macro” in the scikit-learn package in Python.

#### average signal to noise ratio

The average signal to noise ratio was calculated within each cell type. Take 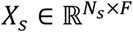 as a subset of raw expression profile *X* ∈ ℝ^*N*×*F*^, where each cell has the same cell type. The average signal to noise ratio is expressed as:

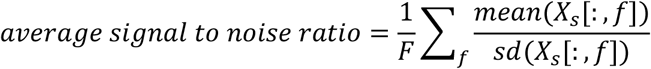

*X_s_*[:, *f*] denotes the column data for protein *f*. *mean*(∙) and *sd*(∙) represent the mean value and the standard deviation. For other methods, *X_s_* in this calculation process will be replaced by the processed expression matrix or embedding matrix.

#### shd

In addition, we defined a spatial heterogeneity degree (*shd*) based on the built cell graph to quantify the spatial heterogeneity. We first compared the cluster assignment of each cell with its spatial neighbouring cells, which were identified by the constructed spatial informative graph. Then the proportion of neighbouring cells with same cluster labels was used as the measure and can be expressed as:

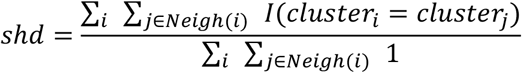

where *j* ∈ *Neigh*(*i*) *if e*_*ij*_ ∈ *E* and *I*(∙) denotes the indicator function. A high *shd* denotes a highly compartmentalized phenotype while a low *shd* denotes a high level of inter-cell type mixing.

## Statistical analysis

The two-sided Mann–Whitney test was utilized for the significance test and was implemented via scipy.stats.mannwhitneyu package.

## Data Availability

All data used in this study are publicly available and the usages were fully illustrated in the Method section.

## Code Availability

The codes are implemented in Python and released on https://github.com/TencentAILabHealthcare/scPROTEIN with detailed instruction.

## Acknowledgements

The authors thank Prof. Ruedi Aebersold for his valuable suggestion for this work. The authors thank Dr. Peilin Zhao for the advice of model development.

## Author Contributions

F.Y. and J.Y. conceived and designed the project. W.L. and H.Z. developed the method. W.L. performed research and conducted experiments under the supervision of F.Y., H.Z. and J.Y.. W.L. and F.Y. analysed the results. W.L. and F.Y. write the manuscript. W.L. finish the figure under the guidance of F.Y. and J.Y.. F.W. helped to polish the figure and manuscript. H.Z. and J.Y. revised the manuscript. Y.R. gave suggestion for the graph model building and the improvement of the manuscript. B.W. gave suggestion for trustworthy AI and the improvement of the manuscript. All authors reviewed and approved the manuscript.

---

**Extended Data Figure 1.**
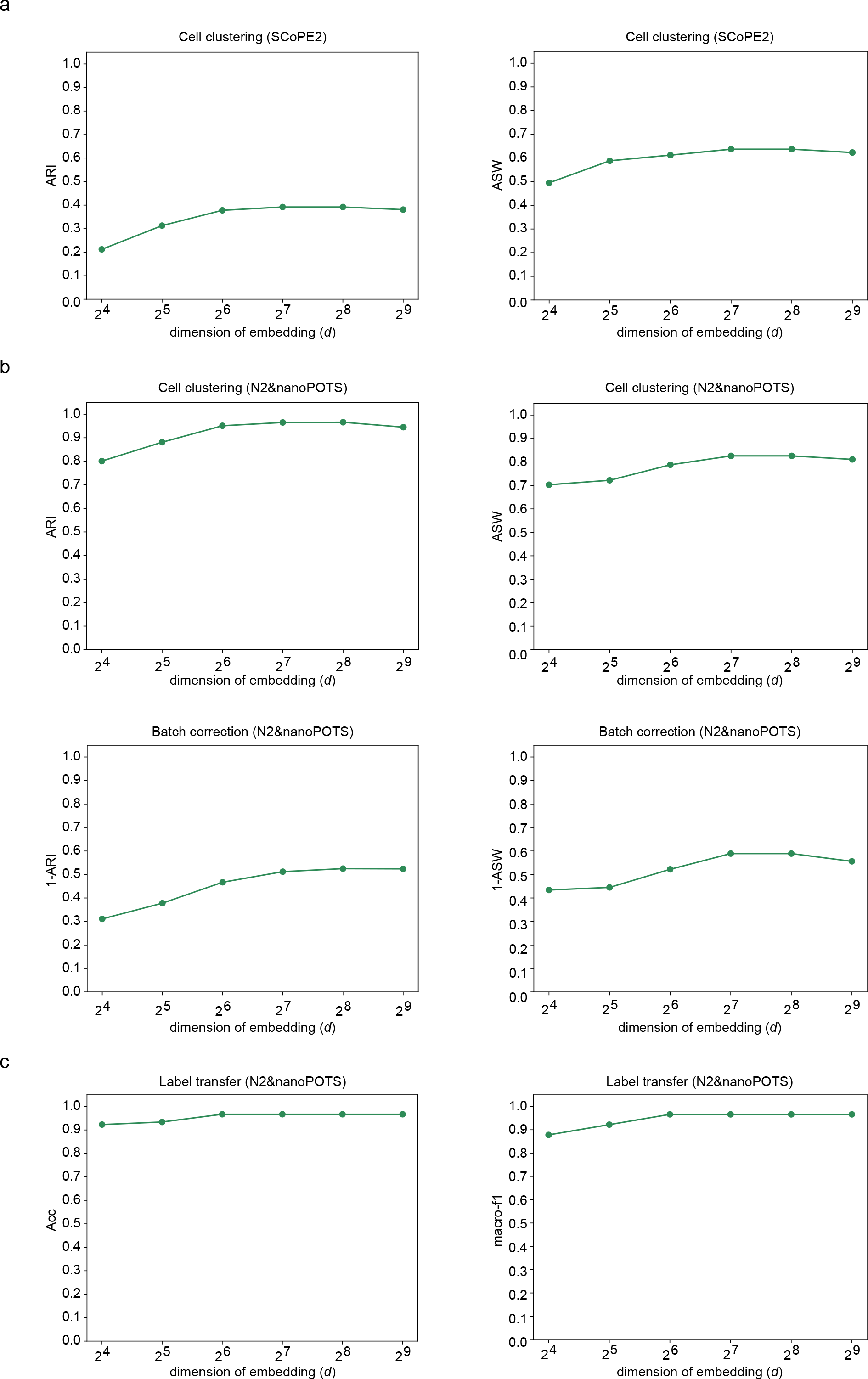
Systematical analysis on the sensitivity of hyperparameter *d*. **a**, The influence of hyperparameter *d* on ASW and ARI in the clustering task using SCoPE2_Specht dataset. *d* is the dimension of learned graph latent embedding of our proposed method. **b**, The influence of hyperparameter *d* on ASW-cell and ARI-cell with cell type labels as ground truth in the batch effect correction task using N2 and nanoPOTS dataset (top panel). The influence of hyperparameters *d* on 1-ASW-batch and 1-ARI-batch with batch labels as ground truth in the batch effect correction task using N2 and nanoPOTS dataset (bottom panel). **c**, The influence of hyperparameter *d* on accuracy and macro-f1 score in the label transfer task (transfer from N2 to nanoPOTS).

**Extended Data Figure 2.**
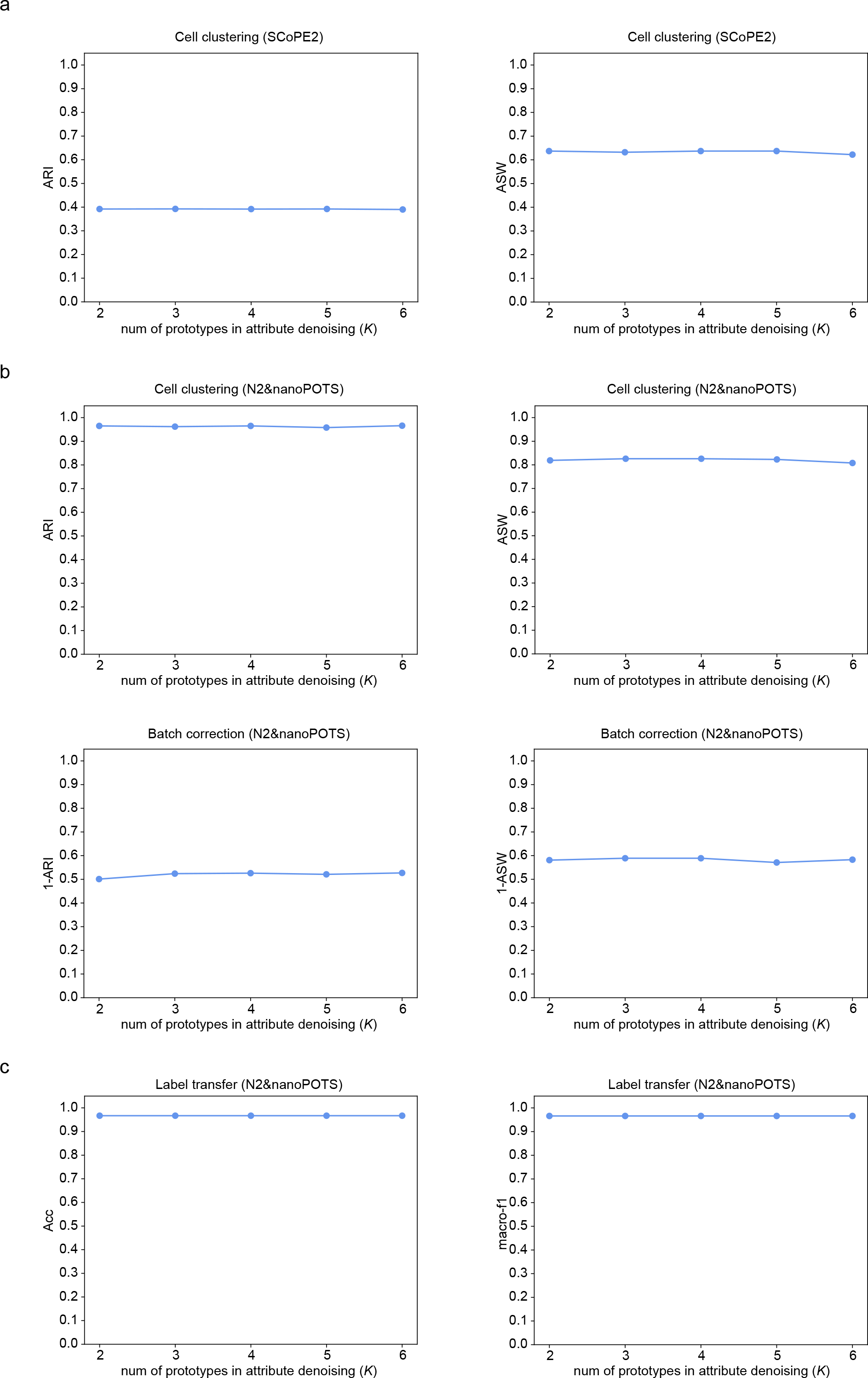
Systematical analysis on the sensitivity of hyperparameter *K*. **a**, The influence of hyperparameter *K* on ASW and ARI in the clustering task using SCoPE2_Specht dataset. *K* is the number of prototypes in attribute denoising of *stage* 2. **b**, The influence of hyperparameter *K* on ASW-cell and ARI-cell with cell type labels as ground truth in the batch effect correction task using N2 and nanoPOTS dataset (top panel). The influence of hyperparameters *K* on 1-ASW-batch and 1-ARI-batch with batch labels as ground truth in the batch effect correction task using N2 and nanoPOTS dataset (bottom panel). **c**, The influence of hyperparameter *K* on accuracy and macro-f1 score in the label transfer task (transfer from N2 to nanoPOTS).

**Extended Data Figure 3.**
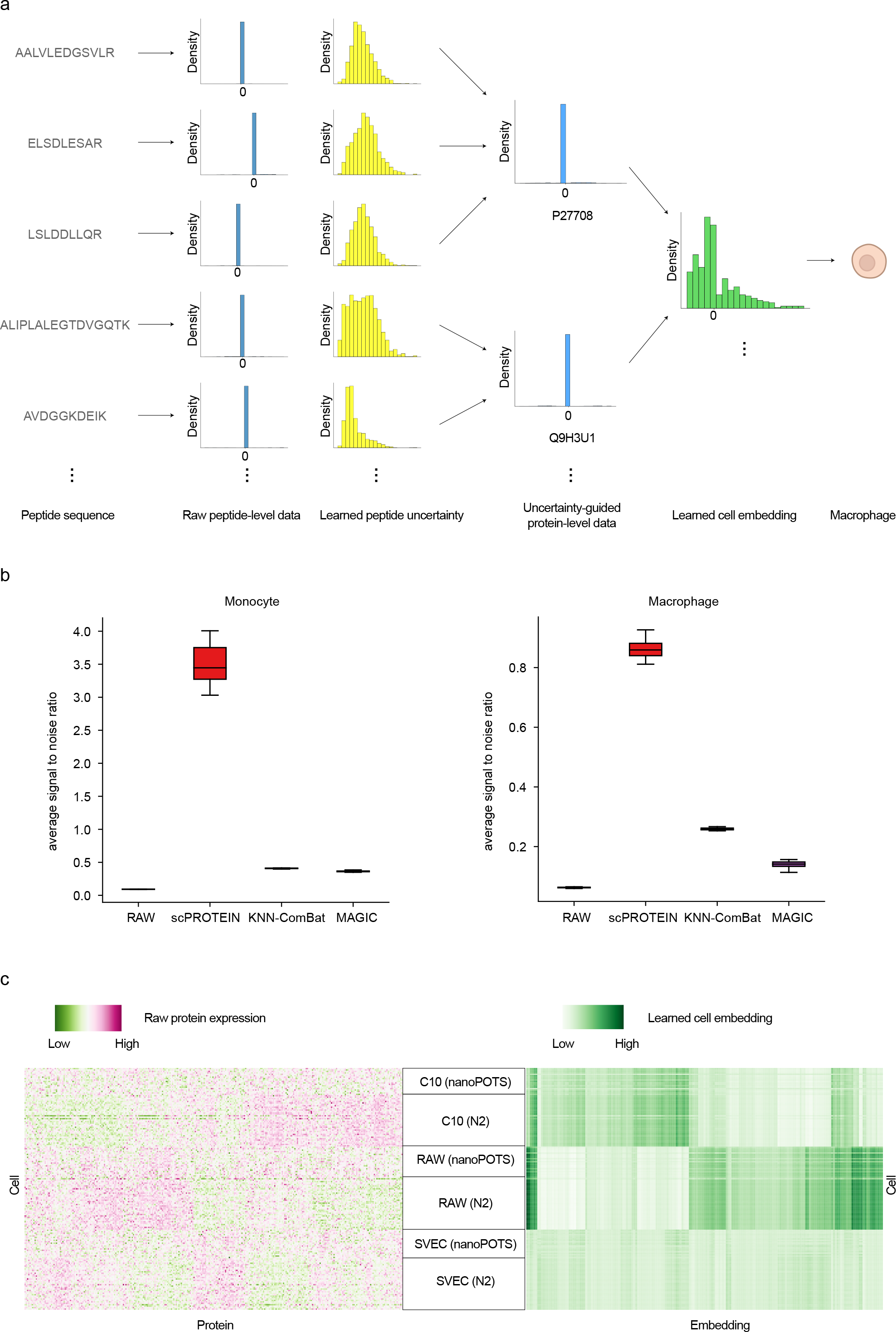
Embedding visualization for SCoPE2_Specht, N2 and nanoPOTS. **a**, The embedding learning process in SCoPE2_Specht dataset. From left to right, we depict the learning process from peptide sequence, raw peptide level data, learned peptide uncertainty for corresponding peptide, aggregated protein level data after uncertainty adjustment and the final learned cell embedding. **b**, The average signal to noise ratio within each cell type of scPROTEIN and comparison methods in SCoPE2_Specht dataset. Higher value indicates better data quality. **c**, The visualization for part of the raw protein expression profile and learned embedding by scPROTEIN in N2 and nanoPOTS datasets. In the left panel, we can observe that the batch effect exhibits in the same cell type across two datasets. In the right panel, scPROTEIN can greatly mitigate the batch effect and the same cell type tend to show similar patterns.

**Extended Data Figure 4.**
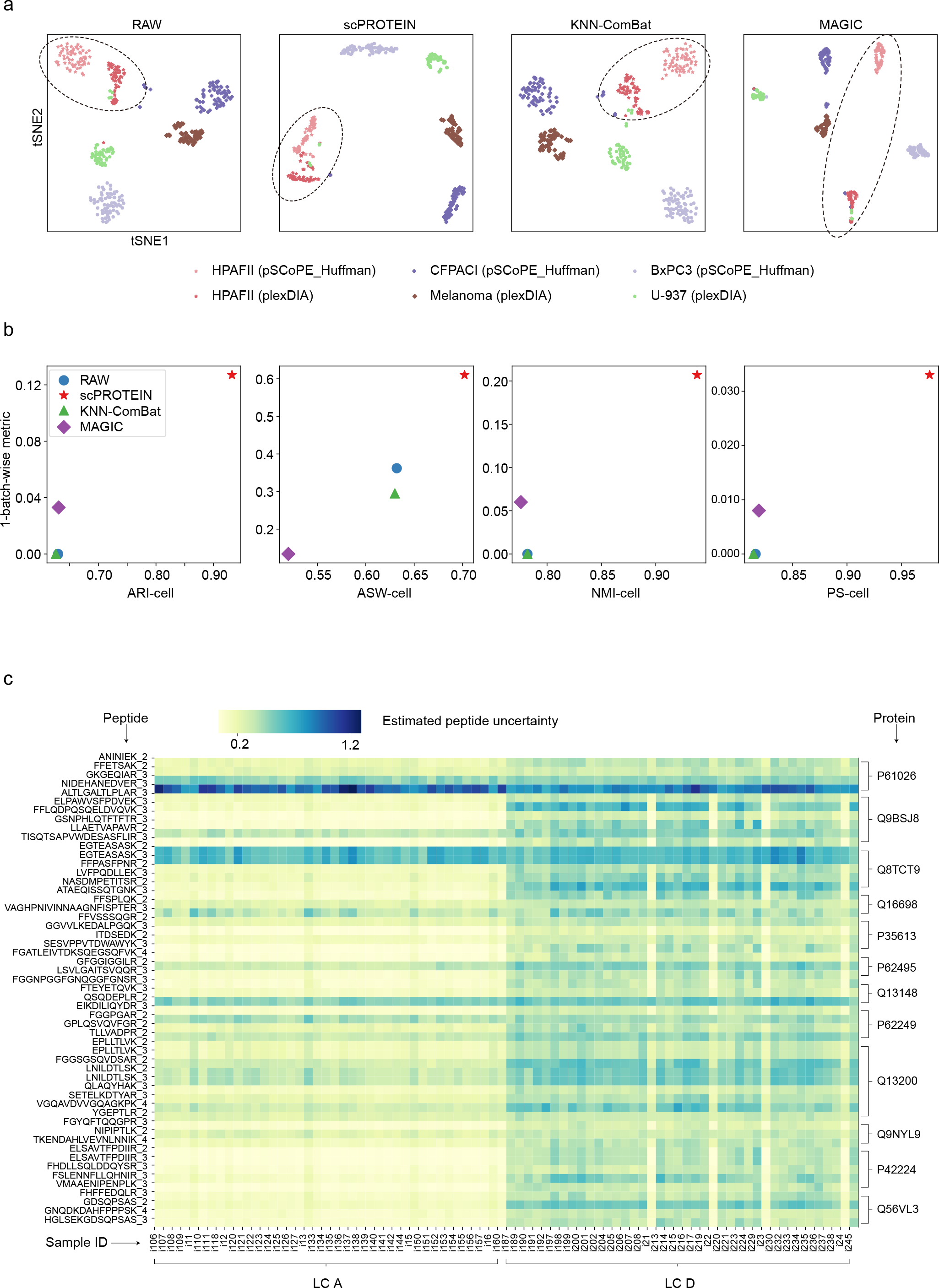
The batch effect correction on cell lines across different datasets. **a**, The tSNE plot showing cells of pSCoPE_Huffman and plexDIA datasets, colored by data source and cell lines. HPAFII is the shared cell line in the two datasets. **b**, The ARI-cell, ASW-cell, NMI-cell, and PS-cell of our proposed method and comparison methods (RAW, KNN-ComBat, and MAGIC) with cell line labels as ground truth (x-axis) and 1-metrics of our proposed method and comparison methods (RAW, KNN-ComBat, and MAGIC) with batch labels as ground truth (y-axis) on pSCoPE_Huffman and plexDIA datasets. **c**, The heatmap showing the estimated uncertainty of each peptide signal across cells, colored by the estimated uncertainty on pSCoPE_Huffman dataset. The batch information and protein information are shown below the heatmap and on the right hand of the heatmap, respectively.

**Extended Data Figure 5.**
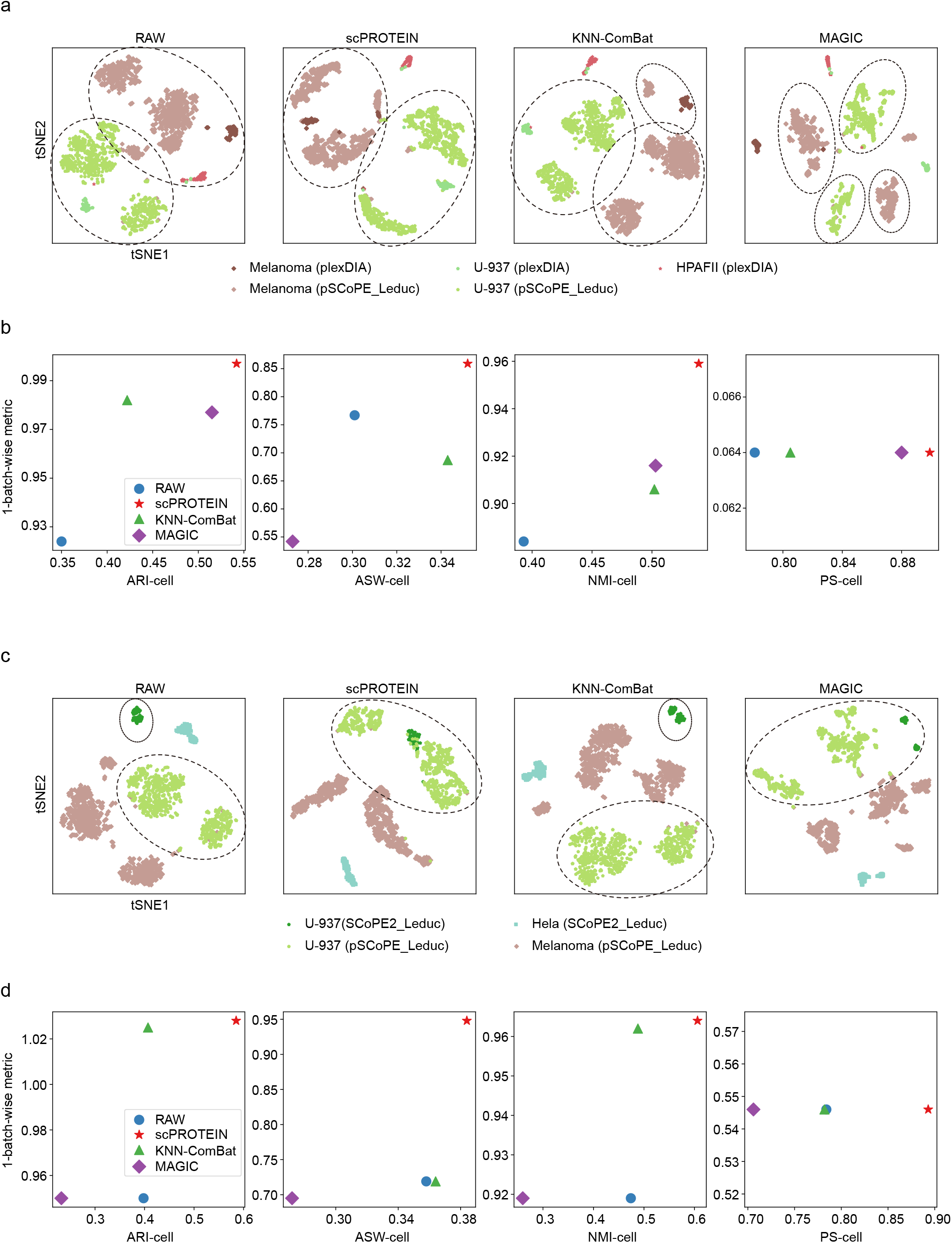
The batch effect correction on cell types across different datasets. **a**, The tSNE plot showing cells of pSCoPE_Leduc and plexDIA datasets, colored by data source and cell lines. Melanoma and U-937 are the shared cell types in the two datasets. **b**, The ARI-cell, ASW-cell, NMI-cell, and PS-cell of our proposed method and comparison methods (RAW, KNN-ComBat, and MAGIC) with cell line labels as ground truth (x-axis) and 1-metrics of our proposed method and comparison methods (RAW, KNN-ComBat, and MAGIC) with batch labels as ground truth (y-axis) on pSCoPE_Leduc and plexDIA datasets. **c**, The tSNE plot showing cells of pSCoPE_Leduc and SCoPE2_Leduc datasets, colored by data source and cell lines. U-937 is the shared cell type in the two datasets. **d**, The ARI-cell, ASW-cell, NMI-cell, and PS-cell of our proposed method and comparison methods (RAW, KNN-ComBat, and MAGIC) with cell line labels as ground truth (x-axis) and 1-metrics of our proposed method and comparison methods (RAW, KNN-ComBat, and MAGIC) with batch labels as ground truth (y-axis) on pSCoPE_Leduc and SCoPE2_Leduc datasets.

**Extended Data Figure 6.**
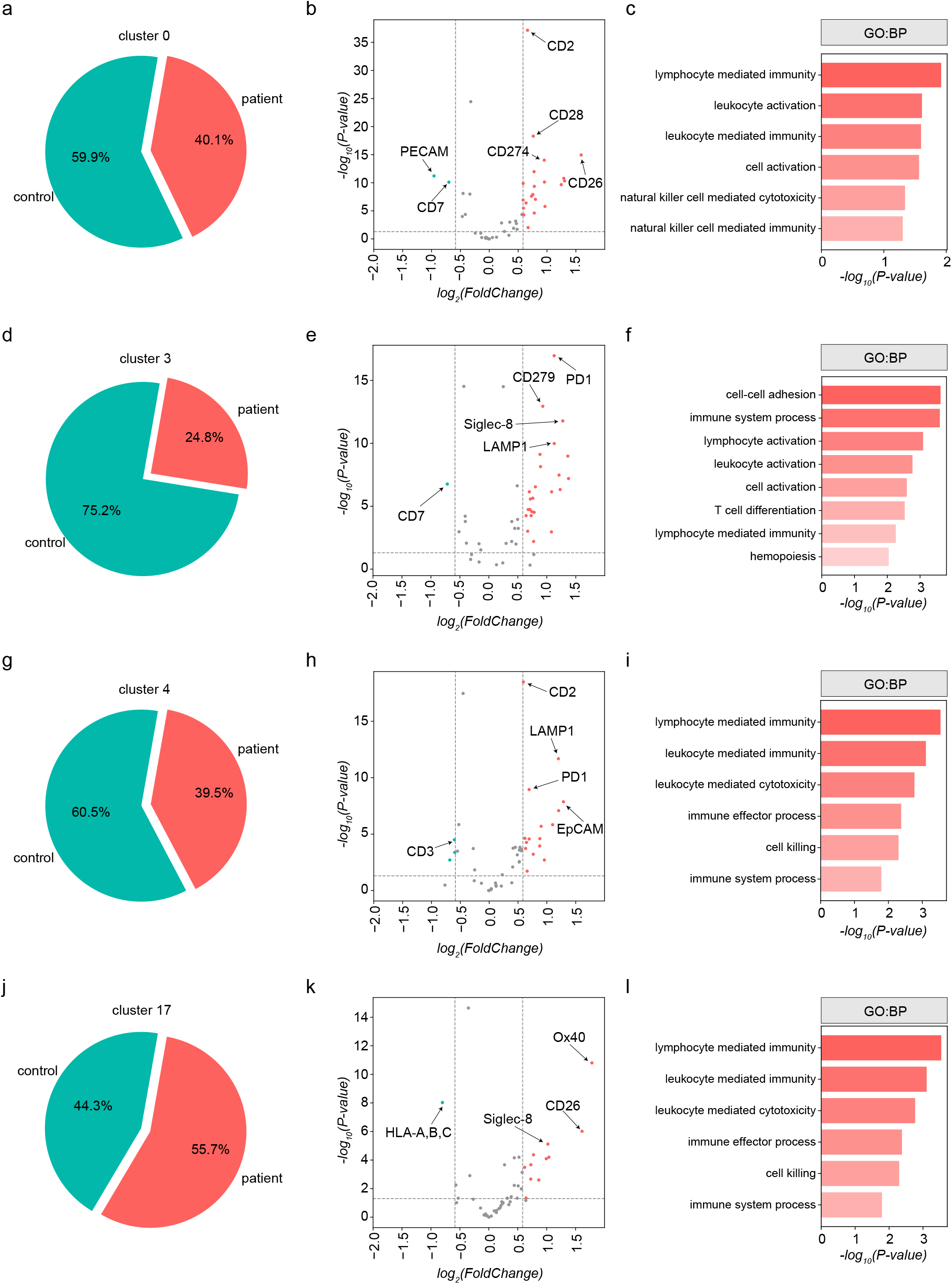
Application of learned embedding on clinical proteomic dataset. **a**, The detailed ratio of cluster 0 cells in control donor and CTCL donor. **b**, The volcano plot showing the differential expression proteins found by contrasting healthy cells and CTCL cells in cluster 0. **c**, Top GO terms in Biological Process (BP) for identified up-regulated proteins for CTCL cells in cluster 0. **d**, The detailed ratio of cluster 3 cells in control donor and CTCL donor. **e**, The volcano plot showing the differential expression proteins found by contrasting healthy cells and CTCL cells in cluster 3. **f**, Top GO terms in Biological Process (BP) for identified up-regulated proteins for CTCL cells in cluster 3. **g**, The detailed ratio of cluster 4 cells in control donor and CTCL donor. **h**, The volcano plot showing the differential expression proteins found by contrasting healthy cells and CTCL cells in cluster 4. **i**, Top GO terms in Biological Process (BP) for identified up-regulated proteins for CTCL cells in cluster 4. **j**, The detailed ratio of cluster 17 cells in control donor and CTCL donor. **k**, The volcano plot showing the differential expression proteins found by contrasting healthy cells and CTCL cells in cluster 17. **l**, Top GO terms in Biological Process (BP) for identified up-regulated proteins for CTCL cells in cluster 17.

**Extended Data Figure 7.**
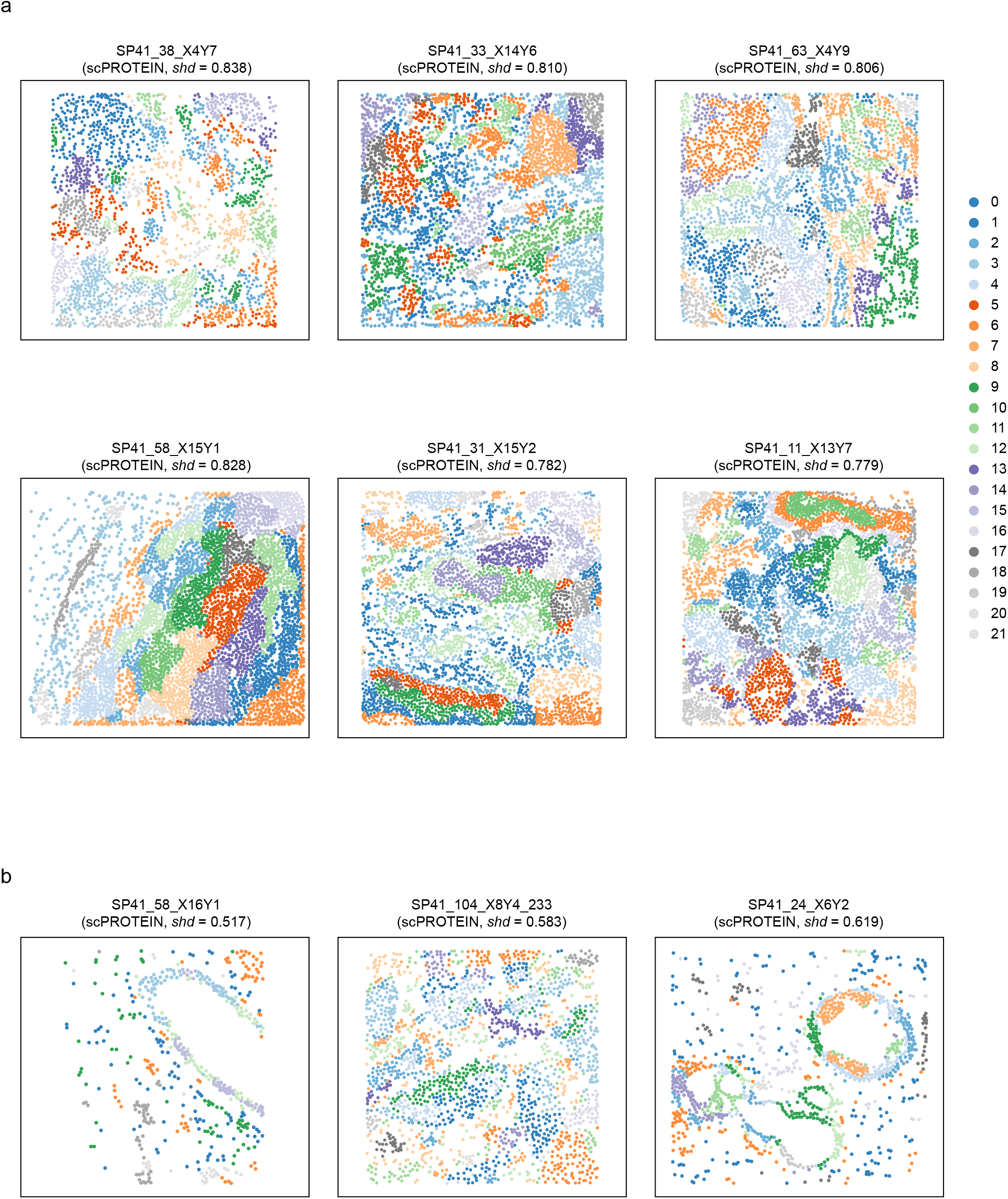
Application of scPROTEIN on spatial proteomic data. **a**, The visualization of clusters on learned spatial informative embedding and the estimated spatial heterogeneity degree within tumor samples. **b**, The visualization of clusters on learned spatial informative embedding and the estimated spatial heterogeneity degree within non-tumor samples.

